# Novel intramolecular base-pairing of the U8 snoRNA underlies a Mendelian form of cerebral small vessel disease

**DOI:** 10.1101/2019.12.12.874594

**Authors:** Andrew P. Badrock, Carolina Uggenti, Ludivine Wacheul, Siobhan Crilly, Emma M. Jenkinson, Gillian I. Rice, Paul R. Kasher, Denis L.J. Lafontaine, Yanick J. Crow, Raymond T. O’Keefe

## Abstract

How mutations in the non-coding U8 snoRNA cause the neurological disorder leukoencephalopathy with calcification and cysts (LCC) is poorly understood. We report the first vertebrate mutant U8 animal model for interrogating LCC-associated pathology. Mutant U8 zebrafish exhibit defective central nervous system development and ribosomal RNA (rRNA) biogenesis, with tp53 activation which monitors ribosome biogenesis. Importantly, LCC patient fibroblasts demonstrate rRNA processing defects. Human precursor-U8 (pre-U8) containing a 3’ extension rescued mutant U8 zebrafish, indicating conserved biological function. Analysis of LCC-associated U8 alleles in zebrafish revealed that one null and one hypomorphic, but still functional, allele combine to cause LCC. Mutations involving any one of seven nucleotides within the human pre-U8 3’ extension, or 5’ region of U8, alter processing of pre-U8, and identify a novel base-pairing interaction between the 5’ end and 3’ extension of human pre-U8. Variants in these seven nucleotides, one of which is present on a single allele in almost all patients, act as hypomorphic mutations. Given that biallelic null U8 alleles are likely incompatible with human development, identification of hypomorphic mutations mediating viable embryogenesis furthers understanding of LCC molecular pathology and cerebral vascular homeostasis.

## Introduction

Leukoencephalopathy with calcifications and cysts (LCC), or Labrune syndrome, is a Mendelian neurological disorder of the cerebral small blood vessels associated with increased morbidity and early mortality, presenting at any age from early infancy to late adulthood. Characterized by the radiological triad of cerebral white matter disease, intracranial calcifications and cysts, LCC was recently shown to be an autosomal recessive genetic disorder caused by biallelic mutations in the gene *SNORD118*, encoding the box C/D U8 small nucleolar RNA (snoRNA) (1).

Ribosomes, the apparatus of protein synthesis, consist of 28S, 18S, 5.8S and 5S ribosomal RNA (rRNA) and 80 core ribosomal proteins distributed into 40S and 60S subunits (2). SnoRNAs are an evolutionarily conserved group of non-protein coding RNAs required for the modification and processing of rRNA. U8, a vertebrate-specific factor, is the only known snoRNA essential for maturation of the 28S and 5.8S rRNAs, components of the 60S large subunit (3). U8 snoRNA is required for removal of the 3’ external transcribed spacer (3’-ETS) sequence, the first of a series of cleavage steps that liberates the 28S, 5.8S and 18S rRNA sequences from the polycistronic precursor-rRNA (pre-rRNA) (4). Pre-U8 snoRNA contains an m^7^G cap and short 3’ extension. Hypermethylation of the m^7^G cap to m3G and removal of the 3’ extension, through a series of steps that appear to involve nucleo-cytoplasmic shuttling and concurrent ordered association and dissociation of multiple protein components (including the LSm proteins, see below), results in the production of mature U8 snoRNA (5–8). The box C/D motif of U8 is bound by four core proteins: 15.5K, NOP56, NOP58 and fibrillarin, contributing to the formation of the U8 small nucleolar ribonucleoprotein (U8 snoRNP) complex and its localization to the nucleolus, the site of pre-rRNA processing (9). The role of U8 in rRNA maturation implicates LCC as a recently discovered novel ribosomopathy, a group of disorders caused by ribosome biogenesis dysfunction that manifest as a diverse set of highly stereotyped clinical syndromes (10).

Of 33 mutation-positive families identified by Jenkinson et al., 31 probands with LCC were compound heterozygotes for two different *SNORD118* variants, implying the existence of one severe (null), and one milder (hypomorphic) mutation, with biallelic null mutations likely incompatible with life. In total, Jenkinson et al., recorded seven putatively causal mutations in the invariant box C/D motif, three within the stem of a conserved hairpin loop which would be predicted to decrease the stability of this structure, three within the highly conserved GAUU motif of the LSm-binding site, and four mutations in the short 3’ extension of the precursor (1). Promoter mutations were also found that reduced expression levels of U8. Presumably, LCC-associated variant combinations reduce U8 function below a critical level while allowing for viable embryogenesis, thus maintaining sufficient levels of production of functional ribosomes. However, the precise molecular pathology of LCC remains unknown.

Here we report the first vertebrate mutant model system to study U8 snoRNA function. Zebrafish U8 mutants were found to exhibit defective rRNA biogenesis and activation of the tumor suppressor p53 (tp53), which monitors ribosome biogenesis dysfunction in a regulatory loop known as ‘nucleolar stress surveillance’ (11–13). Functional assessment of LCC disease-associated U8 alleles confirmed the importance of combinatorial null and hypomorphic mutations. We show that the 3’ extension of U8 is critical for U8 biological activity, an observation reflected in the fact that a mutation within the 3’ extension, or in nucleotides predicted to base-pair with the 3’ extension, were recorded in 29 of 33 patients. Assays using HeLa nuclear cell extracts demonstrated that these mutations alter the processing of pre-U8, revealing a novel secondary structure of human pre-U8 snoRNA. Importantly, fibroblasts from patients with LCC also exhibit rRNA processing defects and human pre-U8 snoRNA was found to rescue the zebrafish U8 mutant, indicating conserved biological function. Taken together, these data support the characterization of LCC as a ribosomopathy whose effects are restricted to the cerebral vessels, and of the utility of zebrafish to provide insight into the pathology of human disease and U8 biology.

## Results

### U8-3 is the predominantly expressed zebrafish U8 during embryogenesis

Zebrafish contain five copies of U8 located on chromosome 10; four copies clustered within the intron of the transcript BX324123, and the remaining copy located between the genes *vamp2* and *and3* (Fig. 1A). Quantitative RT-PCR analysis, exploiting the single nucleotide polymorphisms present between the five copies of U8 for specificity, identified minimal maternal deposition of U8 transcripts in zebrafish, with U8-5 most highly deposited (Fig. 1B, see fig. S1 for alignment of zebrafish U8 copies). At 24 hours post fertilization (hpf) U8-3 was the sole U8 species identified in the zebrafish embryo, with only weak expression of the clustered U8-1, U8-2, U8-4 and U8-5 induced at 48hpf (albeit increasing thereafter) (Fig. 1B). These data provided a rationale for targeted disruption of the U8-3 gene locus to interrogate U8 function during early embryogenesis.

**Fig. 1.**
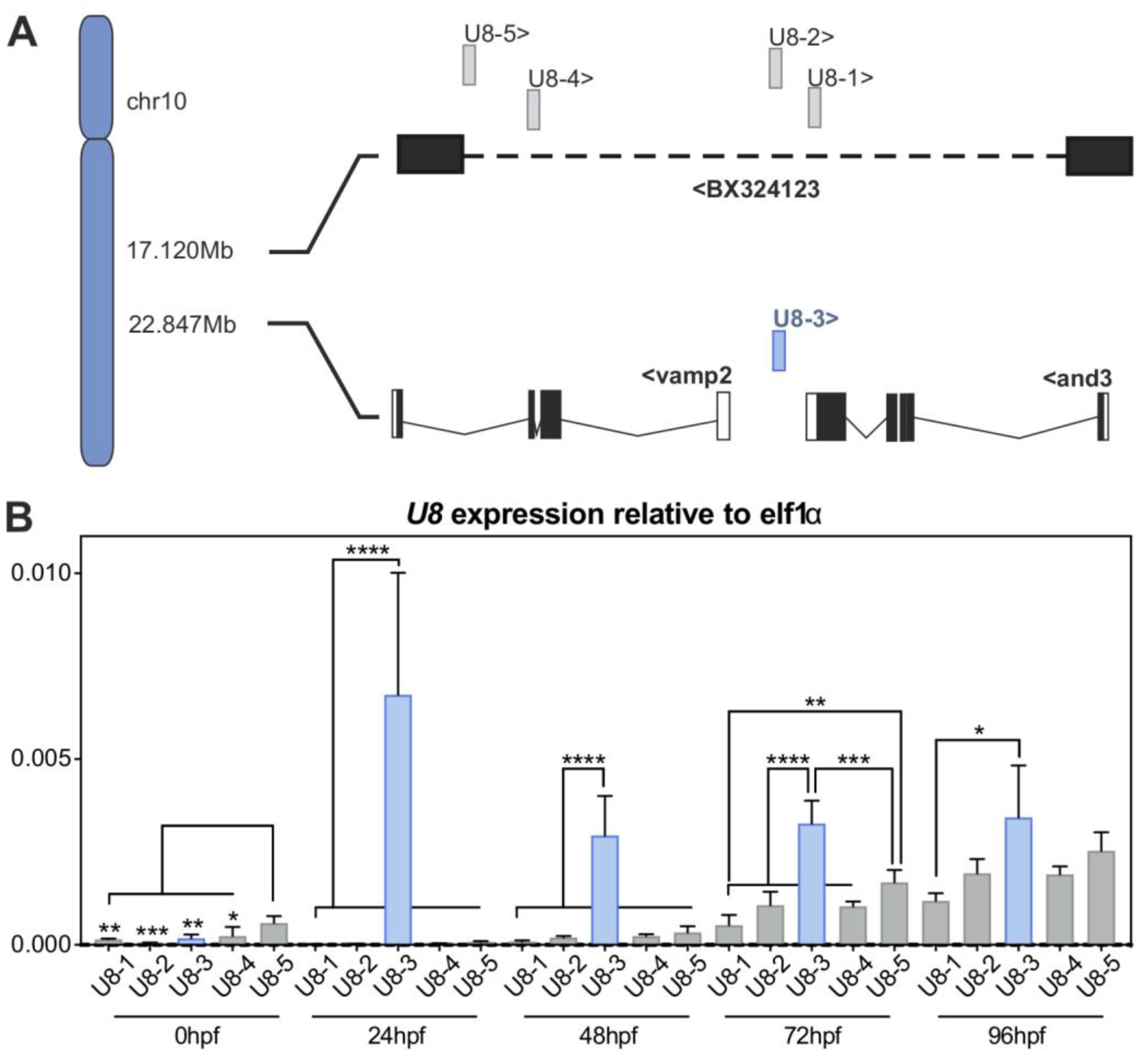
U8-3 is the predominantly expressed U8 snoRNA during zebrafish embryogenesis. **A**) schematic depicting the five U8 zebrafish snoRNA gene copies located on chromosome 10. *U8-3* (blue) is located between the *vamp2* and *and3* genes, whereas the other four copies (grey) are clustered within the intron (dashed line) of the non-coding transcript BX324123. < or > indicates direction of transcription. **B**) quantitative RT-PCR to *U8-1*, *U8-2*, *U8-3*, *U8-4* and *U8-5* snoRNA transcripts at the indicated developmental time points. n=4 biological replicates per time point. Hpf – hours post fertilization.

CRISPR/Cas9 was employed to disrupt the U8-3 locus. Creating insertion/deletion events in a non-coding RNA could lead to unpredictable consequences for U8-3 function. Consequently, two guides were used to excise U8-3 from the genome, producing a null allele, herein referred to as ΔU8-3 (fig. S2). By 24hpf, ΔU8-3 mutants exhibited a less defined midbrain-hindbrain boundary and reduced angiogenic sprouting from the dorsal aorta (Fig. 2A). By 48hpf, these ΔU8-3 mutants demonstrated swelling of the fourth ventricle, consistent with abnormal development of the CNS, reduced melanocyte development, smaller eye size, impaired yolk resorption, disturbed branching of the trunk vasculature and a reduction in embryo length consistent with developmental delay (Fig. 2, A and B). A time course analysis indicated that the ΔU8-3 mutant comes to a developmental standstill, reflected in a failure to resorb yolk and expand the swim bladder (fig. S3). Death was observed from 6 days post fertilization (dpf), with 100% mortality recorded by 9dpf (fig. S4). Quantitative RT-PCR analysis confirmed U8-3 expression is lost in U8-3 mutants (Fig. 2C).

**Fig. 2.**
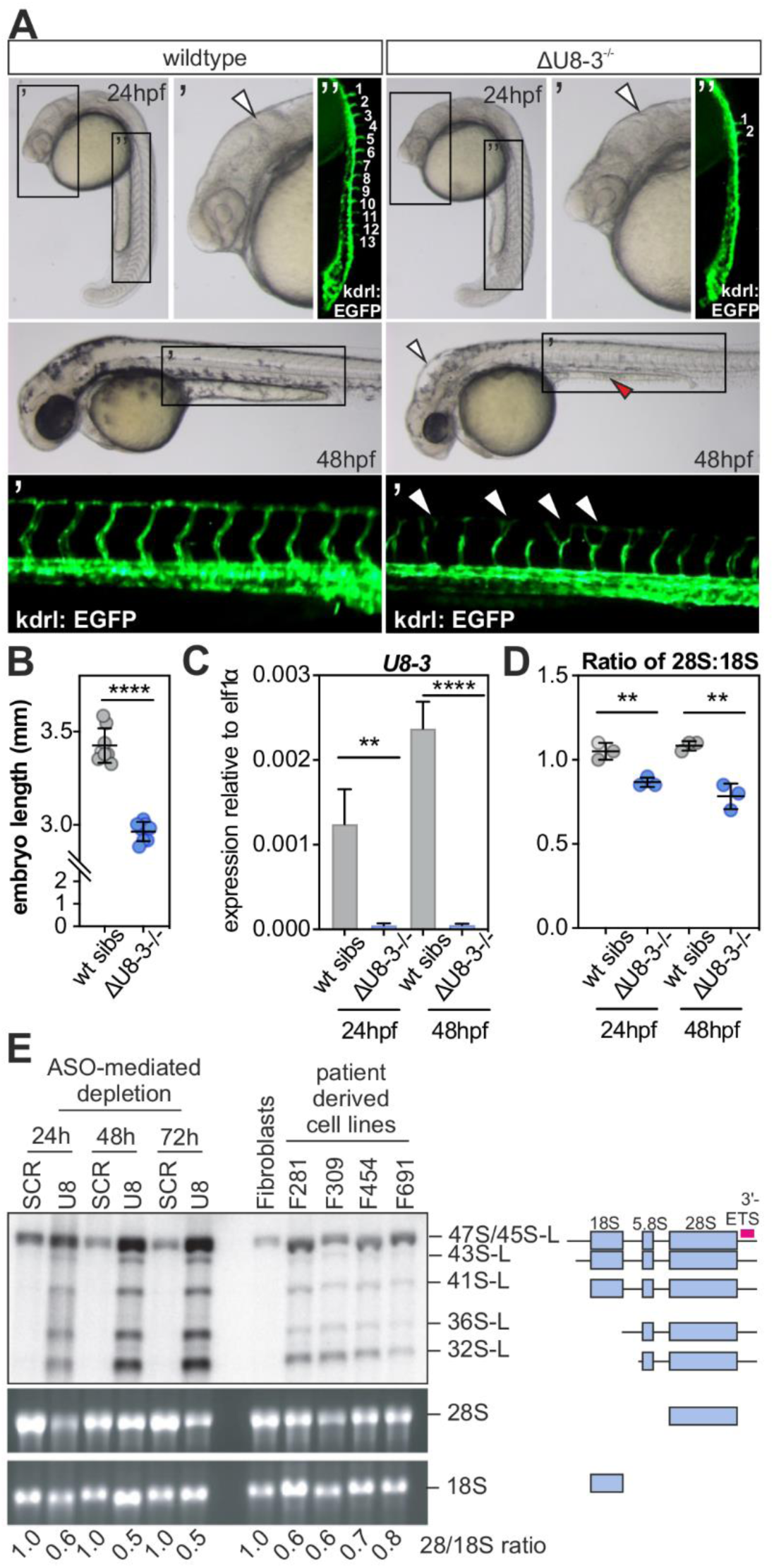
U8-3 is required for correct development of the central nervous system and vasculature in the zebrafish and 28S biogenesis in fish and patient-derived cells. **A**) ΔU8-3 mutants exhibit a less defined midbrain-hindbrain boundary (arrowheads) and reduced angiogenic sprouting (numbered in white, visualized with the kdrl:EGFP transgene) from the dorsal aorta compared to wildtype siblings at 24hpf. By 48hpf ΔU8-3 mutant embryos exhibit hindbrain swelling (arrowhead), an underdeveloped yolk extension (red arrowhead), reduced eye size, impaired melanocyte development and disorganised trunk vasculature compared to wildtype siblings. hpf, hours post fertilization. **B**) quantitation of embryo length of the indicated genotype 48hpf. n=8 embryos per genotype. **C**) qRT-PCR of *U8-3* snoRNA in the indicated genotype and developmental time point. n=4 biological replicates per genotype. **D**) Tapestation analysis demonstrates ΔU8-3 mutant embryos display a preferential reduction in 28S biogenesis consistent with defective 3’-ETS processing. n=3 biological replicates per genotype. **E**) ASO-mediated depletion of human U8 snoRNA results in accumulation of 3’ extended forms of pre-rRNA intermediates (identified by a 3’-ETS specific probe, pink bar) which are required for biogenesis of 28S rRNA. Likewise, these same 3’ extended intermediates accumulate in LCC patient fibroblast cell lines F281, F306, F454 and F691 when compared to wildtype fibroblasts (ATCC). ASO, antisense oligonucleotide. SCR, non-targeting ASO control. The mature 28S:18S ratio, established by densitometry, are indicated.

U8 is required for removal of the 3’ external transcribed spacer (3’-ETS) sequence, and in particular the biogenesis of 28S and 5.8S rRNAs (14–16). Tapestation analysis indicated that U8-3 mutants exhibit a preferential reduction in 28S biogenesis compared to 18S (Fig. 2D). Significantly, northern blotting with a probe specific to the 3’-ETS region of the pre-rRNA demonstrated that LCC patient derived fibroblasts, and control cells in which U8 had been knocked down by antisense oligonucleotides, accumulate aberrant unprocessed 3’ extended rRNA precursors required for 28S biogenesis, which are also impaired for removal of ITS2 (Fig. 2E, L for longform). Further, these pre-rRNA processing defects alter the ratios of 28S to 18S similar to the U8-3 mutant, preferentially impairing biogenesis of 28S (Fig. 2 D, E). Northern blotting analysis with a probe specific to U8 confirmed antisense oligonucleotide-mediated knockdown of U8, and that LCC patient fibroblasts have reduced levels of total U8 most likely due to reduced stability of one, or both, mutant alleles (fig. S5) (1). Taken together, these data support a conserved biological function for U8 in rRNA processing and ribosome biogenesis, and show that patient-derived cells expressing U8 mutations are indeed defective for ITS2 and 3’-ETS maturation.

A portion of the U8-3 promoter was deleted in the ΔU8-3 allele, potentially containing regulatory elements required for the function of other genes. To confirm the specificity of the ΔU8-3 mutant phenotype a complementation test with an independent U8-3 mutant allele was performed. A guide specific to U8-3, the design of which was facilitated by the absence of a relevant PAM sequence in the other zebrafish U8 gene copies, was used to delete 54bp from the U8-3 gene locus, herein referred to as Δ54U8-3 (fig. S6, A and B). Δ54U8-3 mutants demonstrated indistinguishable morphology defects from ΔU8-3 mutants, with both alleles displaying Mendelian autosomal recessive inheritance (fig. S6C). A failure of complementation i.e. the production of 100% wildtype progeny, was observed when a ΔU8-3 heterozygote zebrafish was crossed to a Δ54U8-3 heterozygote zebrafish, demonstrating that the two mutants are associated with loss of function (LOF) of the same gene (fig. S6C).

### Precursor zebrafish U8-3 and human U8 snoRNAs are functionally equivalent

Mutations that lie within the short 3’ extension of human U8 in LCC patients imply that this region is of functional significance (1). However, it has previously been reported that exogenous mature U8 snoRNA, lacking the 3’ extension sequence, localizes to the nucleolus and rescues endogenous U8 depletion in Xenopus oocytes (17). We first performed an electrophoretic mobility shift assay (EMSA) with *in vitro* transcribed mature zebrafish U8-3 and the highly conserved human 15.5K to confirm the ability of *in vitro* synthesized U8-3 to interact with a key U8 snoRNP factor. Addition of 15.5K was found to shift U8-3 migration, and also zebrafish U8-1, 2, 4 and 5, demonstrating that these zebrafish U8 species bind 15.5K (fig. S7). Alignment of human *SNORD118* and zebrafish *U8-3* gene loci identified a putative 3’ extension and 3’ BOX in zebrafish U8 (fig. S8). A transient rescue assay was performed, comparing the capacity of exogenous *in vitro* transcribed mature U8-3 and the putative pre-U8-3 snoRNA to rescue the gross morphological abnormalities observed in the ΔU8-3 mutant. Mature or pre-U8-3 snoRNAs were co-injected into the yolk of 1-cell stage zebrafish ΔU8-3 mutants or wildtype siblings with an mRNA encoding a fluorescent protein acting as a tracer, enabling ubiquitous expression of the transcripts throughout the embryo over the first two days of development (Fig. 3A). Mature U8-3 snoRNA rescued the yolk extension and hindbrain swelling of the ΔU8-3 mutant, but not the effect on embryo length (Fig. 3, B and C). The addition of 14 nucleotides 3’ of the mature U8-3 sequence resulted in a rescue of the hindbrain swelling, yolk extension and embryo length of ΔU83 mutants, thereby demonstrating the biological importance of this 3’ extension sequence (Fig. 3).

**Fig. 3.**
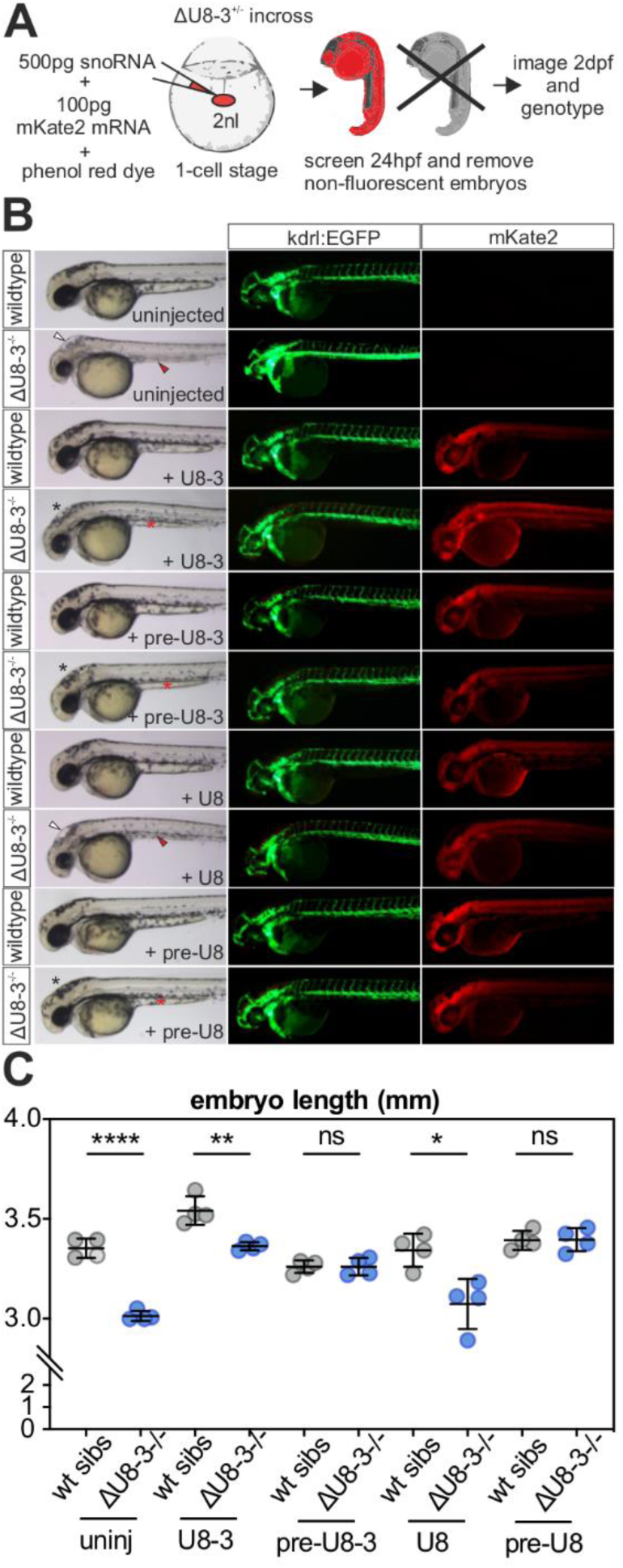
The 3’ extension of precursor U8 is required for optimal U8 biological activity. **A**) schematic depicting the experimental design for transient ubiquitous expression of U8 snoRNA variants and mKate2 mRNA. Bolus size is monitored by a phenol red dye and successful uptake of the microinjected solution into the animal pole traced by red fluorescence. **B**) representative brightfield and fluorescent images of the genotype, exogenous snoRNA, transgenic or fluorescent protein indicated, taken at 48hpf. U8 denotes human sequence whereas U8-3 denotes zebrafish. White arrowheads denotes hindbrain swelling, red arrowheads denotes aberrant yolk extension, black asterisks denotes rescued hindbrain and red asterisks denotes rescued yolk extension. **C**) quantitation of embryo length for the genotypes and introduced snoRNAs indicated. uninj, uninjected. n=4 embryos per genotype.

The above rescue assay provided the means to examine whether human U8 snoRNA is functional in the zebrafish. As for zebrafish pre-U8-3, pre-U8 human snoRNA rescued ΔU8-3 associated gross morphological abnormalities, whereas mature human U8 snoRNA failed to do so (Fig. 3B), indicating that zebrafish and human U8 snoRNA are functionally equivalent. However, although a number of the ΔU8-3 mutant features were rescued by exogenous pre-U8, the eye and head size did not recover to wildtype sibling levels. This lack of complete rescue likely reflects the technical challenge of introducing enough non-coding RNA at the 1-cell stage to sustain a rapidly growing zebrafish embryo over two days of development.

### Functional testing identifies one null and one hypomorphic allele in LCC patients

Having demonstrated that human pre-U8 snoRNA is functional in the zebrafish, we next wanted to test the effect of LCC disease associated mutations in U8-3 null embryos. Six alleles were chosen where a defect in functionality had previously been shown *in vitro*: specifically, either a complete (n57G>A and n58A>G) or reduced (n61A>G) ability to interact with 15.5K compared to wildtype, or disrupted 3’ end processing in HeLa nuclear extracts (n*1C>T, n*5C>G and n*9C>T) (1). Despite molecular evidence suggesting that the n61A>G variant might be hypomorphic, in each case variants affecting nucleotides required for binding to 15.5K were found to act as functional null alleles in that they failed to alter the Δ54U8-3 mutant phenotype, demonstrating the essential nature of this domain for U8 function (Fig. 4). In contrast, all of the patient mutations in the 3’ extension rescued the morphological abnormalities observed in the Δ54U8-3 mutant, including the embryo length defect (Fig. 4 B and C), indicating that their function in ribosome biogenesis is preserved. These data likely reflect the null and hypomorphic status of distinct alleles.

**Fig. 4.**
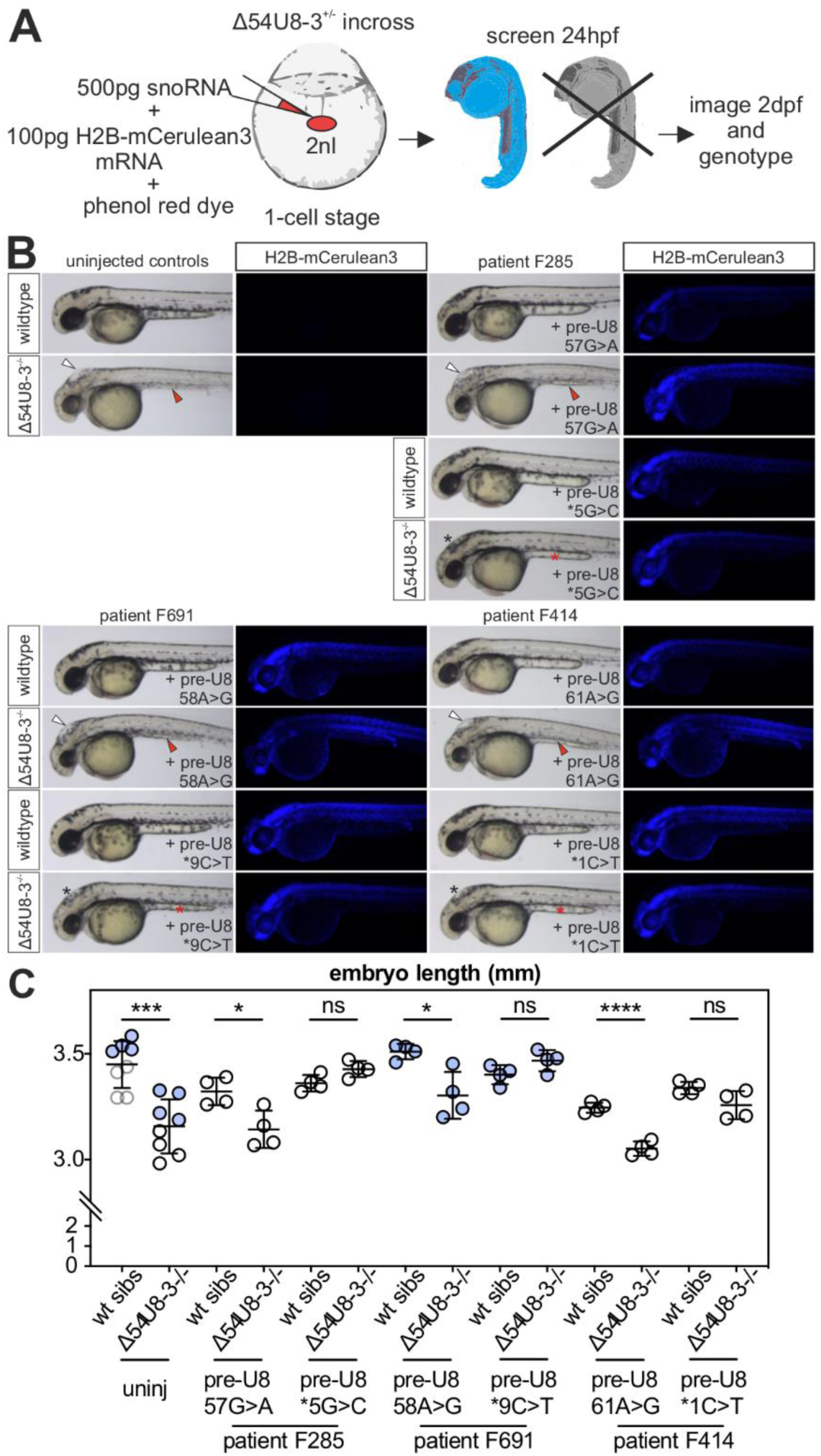
Functional testing of LCC patient mutant U8 snoRNAs identifies one null and one functional allele. **A**) schematic depicting the experimental design for transient ubiquitous expression of U8 snoRNA mutants and H2B-mCerulean3 mRNA. Bolus size is monitored by phenol red dye and successful uptake of the microinjected solution into the animal pole traced by fluorescence. **B**) LCC mutant snoRNAs specific to the region required for 15.5K binding (n57, n58 and n61) fail to rescue the hindbrain swelling (white arrowheads) and yolk extension (red arrowheads) of Δ54U8-3 mutants, whereas LCC mutants specific to the 3’ extension (n*1, n*5, n*9) rescue hindbrain swelling (black asterisks), yolk abnormalities (red asterisks) and pigmentation defects. **C**) LCC mutant snoRNAs specific to the region required for 15.5K binding (n57, n58 and n61) do not rescue the reduced embryo length of Δ54U8-3 mutants, whereas LCC mutants specific to the 3’ extension restore Δ54U8-3 mutant embryo length to that of wildtype siblings. n=4-8 embryos per genotype. Blue and black data points represent embryos collected from two different pairs of heterozygote Δ54U8-3 adults.

### Seven distinct hypomorphic U8 alleles are present in most LCC patients

Two thirds of LCC probands in our cohort have been observed to harbor one of four mutations located within the 3’ extension of pre-U8 (n*1C>T, n*5C>G, n*9C>T, n*10G>T). This observation, and the ability of these mutations to rescue Δ54U8-3 mutant zebrafish reported in Fig. 4, led us to hypothesize that these alleles confer some degree of preserved ribosomal function, and thereby support viable embryogenesis, in the presence of a second, null, mutation on the other allele. Secondary structure predictions of human pre-U8 using RNAfold (18) identified a novel putative base-paring interaction between the 3’ extension of the human pre-U8 snoRNA and the 5’ end of the human pre-U8 snoRNA (Fig 5A). Mapping of all the LCC patient variants on this structure revealed that another three disease-associated-mutations, located at the 5’ end of U8, lie precisely within the proposed base-paired region (Fig. 5A). Strikingly, one of the seven mutated nucleotides found within this novel human pre-U8 base paring interaction was observed in 29 of 33 LCC patients overall (Table S1). Interestingly, this predicted secondary structure of the human pre-U8 snoRNA does not appear to contain the canonical kink-turn found in mature U8 (Fig. 5A). We have previously reported that the n*1C>T, n*5C>G, n*9C>T and n*10G>T mutants demonstrate defective 3’ end processing in HeLa nuclear extracts (1). The predicted secondary structure of human pre-U8, indicating base-pairing between the 5’ end and 3’ extension, combined with the knowledge that n*1C>T, n*5C>G, n*9C>T and n*10G>T mutants cause disrupted processing, suggested that the n2T>C, n3C>T and n8G>C mutations might also affect U8 precursor processing. The human pre-U8 snoRNA is processed to the mature U8 snoRNA in HeLa nuclear extracts after 60 minutes (fig. S9). When examining U8 processing intermediates in HeLa nuclear extracts at 30 minutes, a time point before mature U8 snoRNA is produced, we observed that each of the n2T>C, n3C>T and n8G>C mutants conferred an aberrant, increased, rate of 3’ end processing when compared to wildtype (Fig. 5B).

**Fig. 5.**
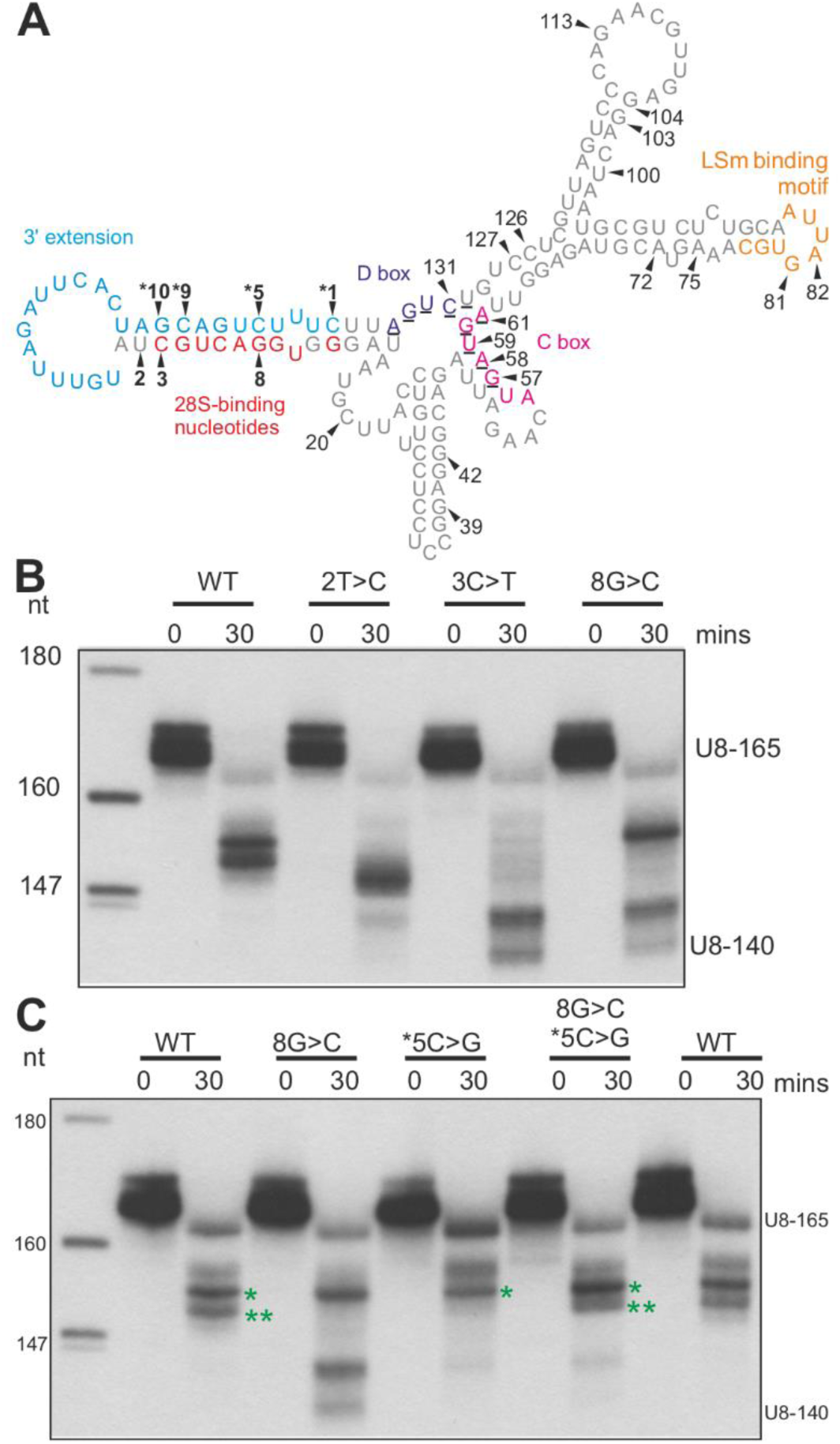
Aberrantly processed mutant U8 snoRNA alleles are present in 29 of 33 LCC patients. **A**) schematic depicting the secondary structure of human precursor U8 predicted by RNAfold software (18). Black arrowheads/bold numbering identify seven mutated nucleotides in LCC patients that lie within a base-pairing region encompassing the 3’ extension. The remaining mutated nucleotides recorded in LCC patients are numbered (refer to Supplementary **Table S1** for a complete list of LCC-associated mutant alleles). The novel 5’ end/3’ extension base-pairing may preclude premature kink-turn formation (normally involving residues: 57-61 and 130-134, underlined) until U8 has matured sufficiently. **B**) processing of 5’ end-radiolabeled *in vitro* transcribed precursor U8 wildtype/mutant snoRNA (U8-165) was assessed in HeLa nuclear extracts. At 30 min, 2T>C, 3C>T, and 8G>C mutant U8 snoRNAs exhibit an enhanced rate of processing when compared to wildtype as demonstrated by the presence of mature U8 (U8-140). **C**) processing of 5’ end-radiolabeled *in vitro* transcribed precursor U8 wildtype/mutant snoRNA (U8-165) was assessed in HeLa nuclear extracts. At 30 min, the n*5C>G mutant U8 is blocked in processing when compared to wildtype (** band is absent), in contrast to 8G>C mutant U8 snoRNA that exhibits production of mature U8 (U8-140). When base-pairing complementarity is restored by combining the two mutations the pattern of processing is restored to wildtype.

If the proposed interaction of the 5’ and 3’ ends of human pre-U8 is correct, then restoring base-pairing complementarity for the n8G>C mutation would be predicted to return processing to wildtype. Thus, the n8G>C and n*5C>G mutations were combined to test this hypothesis. n*5C>G alone appeared to exhibit slowed processing when compared to wildtype at the early 30 minute timepoint, whereas n8G>C was again associated with an increased rate of processing and production of mature U8 (U8-140) after 30 minutes (Fig. 5C). As predicted by the model, the n8G>C/n*5C>G double mutant pre-U8 was found to confer almost identical processing when compared to wildtype, providing functional evidence for the proposed base-pairing between the 5’ end and 3’ extension of the human pre-U8 snoRNA (Fig. 5C).

The very survival of LCC patients beyond embryogenesis suggests that, like the 3’ extension mutations tested in Fig. 4, the n2T>C, n3C>T, n8G>C and n*10G>T mutant pre-U8 snoRNAs also retain some degree of functional competence necessary for ribosome biogenesis. To test this, rescue experiments using the n2T>C, n3C>T, n8G>C and n*10G>T mutants were performed in the zebrafish Δ54U8-3 mutant. Consistent with our hypothesis, n2T>C, n8G>C and n*10G>T pre-U8 all rescued Δ54U8-3 mutant morphology, including embryo length (fig. S10, A and B). In contrast, the n3C>T pre-U8 did not salvage the yolk extension defect, or embryo length (fig. S10, A and B). However, it should be noted that almost half the nucleotides required for 28S binding in human differ from zebrafish (fig. S10C), and such a lack of conservation might adversely affect the capacity of human U8 to completely substitute for U8-3 function in the zebrafish when single base pair changes are introduced within the 5’ end of human U8. As a consequence, caution must be exercised when interpreting a failure to rescue in these circumstances.

### tp53 is activated in response to loss of U8-3 in zebrafish

Perturbation of ribosome biogenesis activates the transcription factor TP53, a tumor suppressor with a role in a wide range of biological processes, including DNA damage, mitochondrial stress, autophagy and oncogenesis (19). Trans-activated tp53 increases expression of different effectors depending on the biological context, including *Δ113tp53*, transcribed from intron 4 of the tp53 gene in zebrafish. *Δ113tp53* was found to be upregulated 50-fold in ΔU8-3 mutants compared to wildtype siblings at 24hpf, as were the tp53 target genes *mdm2, cyclinG1, p21 and bax* (Fig. 6A, fig. S11A). To determine which tissues were affected by loss of *U8-3* in zebrafish, a spatio-temporal reporter of tp53 activity was generated that utilized the *Δ113tp53* promoter containing two tp53 binding sites (Fig. 6B) (20). At 17hpf, prior to any observable morphological abnormalities, tp53 trans-activation activity can be observed already in ΔU8-3 mutants (fig. S11B). By 24hpf, tp53 trans-activation activity was detected in the eye, CNS and somites of ΔU8-3 mutants (Fig. 6C, fig. S11C), and by 48hpf, the CNS and somites were the most highly fluorescent tissues (Fig. S11D). Importantly, somite derived vegf expression is critical for correct patterning of the dorsal aorta and for arterial development (21). As such, the delayed sprouting and abnormal branching of the trunk vasculature is unlikely to be a cell autonomous effect, but rather to be secondary to the impairment of somitogenesis. The tissues in which tp53 is activated all display obviously impaired development in the ΔU8-3 mutant as evidence by the reduced eye size, shortened body length and fourth ventricular swelling (fig. S11D).

**Fig. 6.**
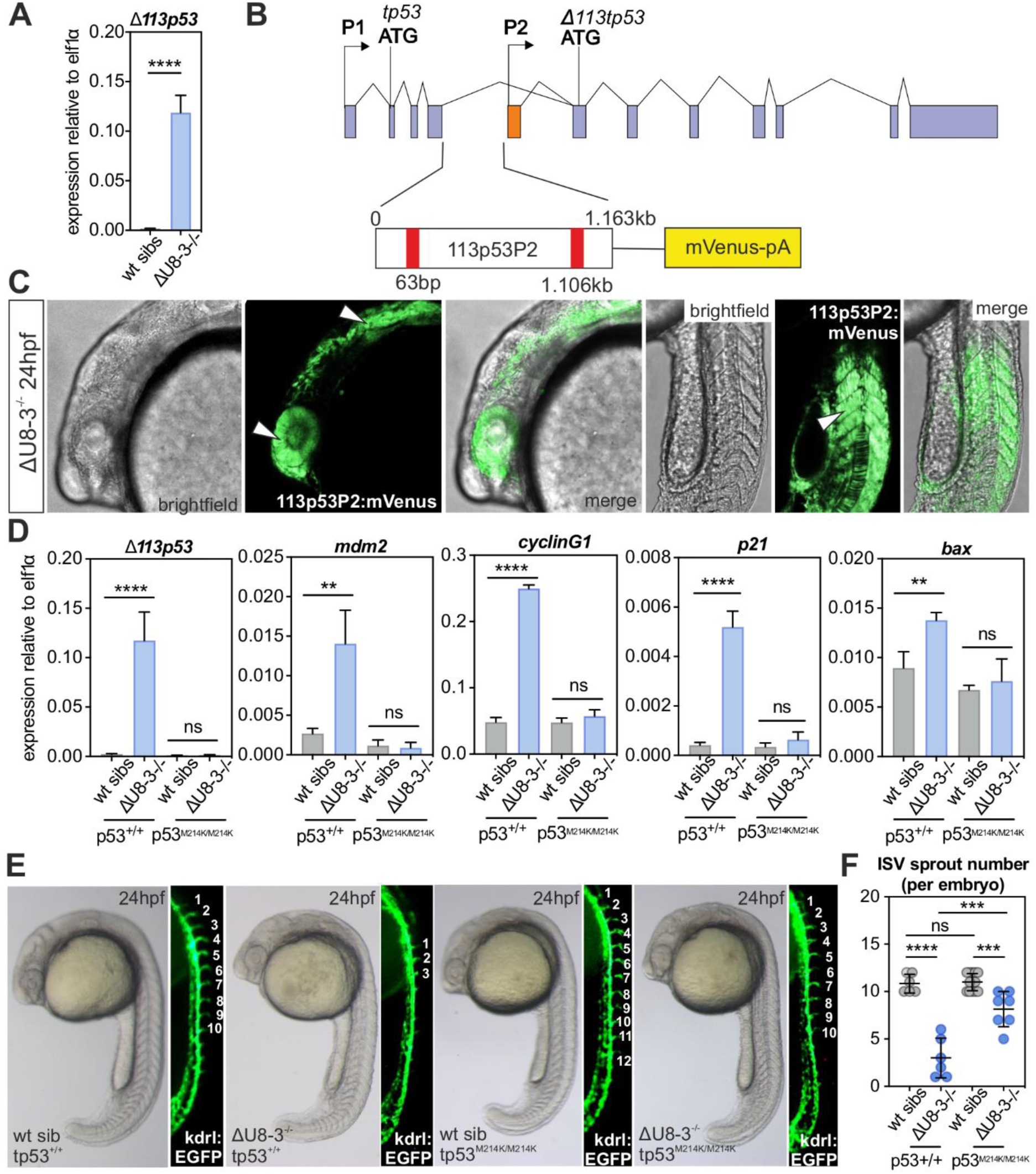
tp53 is trans-activated in response to loss of *U8-3* in zebrafish. **A**) quantitative RT-PCR of *Δ113tp53* transcripts 24hpf. **B**) schematic depicting 1.163kb of intron 4 of the tp53 gene that contains two tp53 binding sites (red bars) used to drive expression of mVenus when tp53 is trans-activated. **C**) tp53 activity is observed in numerous tissues that develop abnormally in ΔU8-3 mutants, including the eye, hind-brain and somites (arrowheads) at 24hpf. **D**) quantitative RT-PCR demonstrates significant upregulation of the tp53 target genes *Δ113tp53, mdm2*, *cyclinG1*, *p21* and *bax* in ΔU8-3 mutants when compared to wildtype siblings at 48hpf, that is abrogated by genetic inactivation of tp53. hpf – hours post fertilization. n=4 biological replicates per genotype. **E**) representative brightfield and corresponding fluorescent images of the indicated genotype and transgenic reporter. Angiogenic sprout number indicated by white numbering. **F**) quantitation of angiogenic sprouts in the indicated genotype. n=6-13 embryos per genotype.

### Inactivation of tp53 signaling partially rescues the ΔU8-3 mutant

To characterize the consequences of tp53 signaling in the ΔU8-3 mutant, ΔU8-3 mutant embryos were crossed onto a tp53 mutant background (22). Notably, genetic inactivation of tp53 prevented induction of *Δ113tp53*, *mdm2*, *cyclinG1*, *p21*, and *bax* expression in ΔU8-3 mutants that is typically observed at 48hpf (Fig. 6D), confirming that the increased expression of these mRNAs is tp53-dependent. Genetic inactivation of tp53 was found to partially restore angiogenic sprouting in ΔU8-3 mutants (Fig. 6E, F), and to rescue ventricular swelling 48hpf (fig. S12A). Although tp53 clearly contributes to the gross morphological abnormalities of ΔU8-3 mutants, inactivation of tp53 signaling would not be predicted to restore the defect in rRNA processing. Indeed, tapestation analysis demonstrated that the preferential inhibition of 28S biogenesis is not rescued in tp53 mutant ΔU8-3 mutant embryos (fig. S12B), and these embryos remain significantly shorter than their U8-3 wildtype sibling counterparts (fig. S12C). Taken together, these data indicate that the reduced embryo length of ΔU8-3 mutants results from impaired ribosome function.

## Discussion

Informed by the genetic architecture of the Mendelian disease LCC, here we describe the use of a novel vertebrate mutant animal model of U8 dysfunction to shed new light on the processing of the snoRNA U8, its role in ribosome biogenesis, and the pathology of LCC.

Our data indicate that both zebrafish U8 mutant animals and fibroblasts from patients with LCC exhibit rRNA processing defects. The majority of human syndromes linked to defective ribosome biogenesis are thought to result from haploinsufficiency or partial loss of function of ribosomal proteins or ribosome assembly factors (23–25), and LCC patient-derived fibroblasts express reduced amounts of U8. In our previously published series, 31 of 33 probands with LCC were compound heterozygous for two different *SNORD118* mutant alleles (1). Given that most rare autosomal recessive traits demonstrate enrichment for consanguinity and allelic homozygosity, these molecular data indicate that biallelic null mutations in *SNORD118* are likely incompatible with development. In keeping with this hypothesis, we show that the U8-3 zebrafish mutant is embryonic lethal. Furthermore, functional rescue assays in mutant zebrafish confirm the importance of combinatorial null and hypomorphic mutations in LCC pathology, with partial loss-of-function alleles mediating viable embryogenesis.

The biogenesis and processing of essential, independently transcribed snoRNAs is dependent on highly organized secondary structures and their sequential association with core snoRNP proteins that halt advancing exonucleases, with nucleolytic trimming of a precursor RNA necessary to achieve functionality and to provide metabolic stability and nucleolar localization (7,8,26). Using our zebrafish model we were able to shed new light on the processing of human U8, identifying a novel base-pairing interaction between the 5’ end and 3’ extension of human pre-U8. Twenty one of 33 patients identified by Jenkinson et al. harbored at least one mutation within the 3’ extension, whilst a further eight patients were positive for a mutation in a 5’ situated nucleotide predicted to base-pair with this 3’ extension (1). Human pre-U8 snoRNA containing a 3’ extension rescued the zebrafish U8 mutant, indicating a conserved biological function. To our knowledge, the requirement for such nucleotide base-pairing within the human pre-U8 snoRNA has not been described previously. Close examination of the five zebrafish pre-U8 snoRNAs does not reveal an extensive potential for base-pairing between the 5’ end and the 3’ extension as proposed for the human pre-U8 snoRNA. Any base-pairing potential of the zebrafish pre-U8 snoRNAs would require exact mapping of the 5’ and 3’ ends of the zebrafish pre-U8 snoRNAs and experimental validation. We hypothesize that in human cells, the base-pairing between the 5’ and 3’ ends of the human pre-U8 snoRNA may be important to regulate the timing of final maturation of U8 by either masking its functionally important 5’ end and/or to prevent premature formation of a canonical kink-turn, and associated protein binding, until the U8 snoRNP has sufficiently matured. Such regulation may be important for the correct processing of U8, its metabolic stability, its nucleo-cytoplasmic trafficking (7, 27), its final targeting to the nucleolus, or even possibly to facilitate extra-ribosomal functions of U8 independent of its interaction with the pre-rRNAs. Previous work from the Lührmann laboratory identified four U8 snoRNA precursor processing intermediates, leading them to hypothesize that either the U8 pre-snoRNA is processed by more than one exonuclease activity, or that the sequence of the 3’ extension may determine the kinetics of 3’ processing (7). Our data, as presented in Figure 5, are in agreement with the latter hypothesis, as LCC-associated single nucleotides changes located within the 3’ extension, or that pair with this region, appear to alter the kinetics of processing of the U8 pre-snoRNA. The assembly, processing and export of small nuclear RNPs (snRNPs) also follows a well-defined pathway like that described for snoRNPs (28–30). Precursor forms of the spliceosomal U2 small nuclear RNA (snRNA) pair the 3’ extension with an internal sequence, this base-pairing being essential for correct processing into a mature snRNP (31). It is possible that base-pairing of 3’ extension sequences may be a more general characteristic of snRNA/snoRNA maturation.

In cell culture models, tp53 is required for activation of cell cycle arrest within 24 hours from the onset of ribosomal stress before, over time, impaired ribosome function becomes rate limiting for cellular growth and division (11,32–34). Depletion of U8 has recently been shown to result in potent induction of TP53 in human cells (13, 16), and tp53 signaling in the U8-3 mutant would be predicted to induce an earlier and more complete inhibition of the cell cycle. In keeping with this, the angiogenic sprouting defect observed in the U8-3 mutant at 24hpf was largely, but not completely, rescued by genetic inactivation of tp53; with the partial rescue probably explained by a preferential reduction of 28S in U8-3 mutants indicating that ribosomal dysfunction is already manifested by this time. By 48hpf, when ribosome dysfunction is likely more pronounced, the morphology of the U8-3 mutant was only minimally salvaged. TP53 is normally constitutively degraded by the proteasome following Hdm2-mediated ubiquitination. When ribosome biogenesis is compromised, unassembled ribosomal components accumulate; this is notably the case for a trimeric complex consisting of the two ribosomal proteins uL5 and uL18 and the 5S rRNA which captures Hdm2, titrating it away from TP53, resulting in a net stabilization of TP53 and the activation of a cell death program (35, 36). Although knockdown of U8 in human cells would, therefore, indirectly activate TP53, it remains to be determined whether such activation occurs in the affected neurological tissue of patients with LCC, and whether this plays a mechanistic role in disease pathology.

Alterations in ribosomal components underlie a heterogeneous class of diseases referred to as the ribosomopathies, with a diversity in associated clinical phenotype providing an indication of the multiple specialized roles of the ribosome in normal physiology. Despite ubiquitous expression of the U8 snoRNA, germline mutations in *SNORD118* cause a progressive microangiopathy apparently limited to the cerebral vasculature. Although the phenotype of the exclusively neurological disease LCC is highly distinctive, it is not pathognomonic, as a remarkably similar radiological association is seen in the context of the multisystem disorder Coats plus. Coats plus is caused by mutations in *CTC1* (37) and *STN1* (38), both components of the conserved heterotrimeric telomeric capping complex, and in the telomeric protein POT1 (39). Interestingly then, an unbiased enChip-RNAseq approach identified U8 as a telomere associated RNA (40). As such, the precise link between U8 and cerebral vascular homeostasis awaits elucidation, and may conceivably involve both ribosomal defects, that activate TP53 and/or induce selective impairment of translation of distinct mRNAs in a cell lineage specific context, or currently undefined non-ribosomal functions of U8.

## Materials and Methods

### Zebrafish strains and husbandry

Establishment and characterization of the *tp53^M214K/M214K^* and *Tg(kdrl:GFP)^s^*^843^ strains have been described elsewhere (22, 41). Embryos and adults were maintained under standard laboratory conditions as described previously (42) and experiments were approved by the University of Manchester Ethical Review Board and performed according to UK Home Office regulations. No statistical method was used to predetermine sample size for experimental groups.

### Real-time quantitative PCR

RNA was isolated from homogenised zebrafish embryos using TRIzol (Thermo Fisher Scientific) and genomic DNA removed with the TURBO DNA-free Kit (Thermo Fisher Scientific). Reverse transcriptase was performed with the ProtoScript II First Strand cDNA synthesis kit (New England BioLabs) using 1µg of total RNA with random hexamer primers. qRT–PCR analysis was performed with the primers described in table S2 using a 60°C annealing temperature, primers with efficiencies between 95-105%, and the SensiFAST SYBR No-ROX kit (Bioline) and the Mx3000P system (Stratagene).

### Genome editing

Capped nls-zCas9-nls, mKate2 or H2B-mCerulean3 mRNA was synthesized using a mMESSAGE mMACHINE SP6 kit (Life Technologies) from a linearized pCS2 construct and purified using a RNeasy mini kit (Qiagen). Zebrafish guides were designed using the Harvard chopchop program (https://chopchop.rc.fas.harvard.edu). Guide RNA (gRNA) incorporating this target sequence was generated from a PCR amplification product (see table S2 for primer sequences) including the remaining sequence of *S. pyogenes* chimeric single guide RNA through *in vitro* transcription using a HiScribe T7 Quick kit (New England Biolabs). The gRNA was then precipitated in a 1/10 volume of 3M Sodium Acetate and two volumes of 100% ethanol by chilling the reaction at −20 °C for 15 min, then spinning in a microcentrifuge (sigma) at 13K for 15 min, and finally the RNA pellet was resuspended in 15 μl of RNase free water. Cas9 mRNA (250 pg) and gRNA (30 pg) and mKate2 (100pg) or H2B-Cerulean3 (100pg) were injected into the yolk of 1-cell stage embryos and fluorescence used to identify successfully injected embryos. Working guides were identified by PCR amplifying the target region and running the PCR product on a 3% agarose gel to identify INDEL events that produce visible shifts or smearing of the amplification product.

### Genotyping

Embryos or fin-clips were placed in PCR tubes with 50 μl of 50 mM NaOH and denatured for 20 min at 95 °C. A volume of 20 μl of Tris-HCl pH 8 was added to each tube and 1 μl of the genomic DNA used for PCR amplification

### Polymerase chain reaction conditions

Polymerase chain reaction (PCR) was performed in a 25–50 μl reaction mix containing DNA template (0.1–100 ng DNA), sense and antisense primer 0.8μm each, 0.25 mM dNTPs (Bioline), 1X HF buffer (New England Biolabs), 1U Phusion Taq polymerase (New England Biolabs), 0.5 mM MgCl (New England Biolabs) and 1.5 μl DMSO (New England Biolabs) per 50 μl reaction. PCR was performed in a Techne TC-PLUS or Alpha Thermal Cycler PCR^MAX^ machine with an initial denaturing step at 98 °C for 3 min followed by 35 cycles of denaturing at 98 °C for 10 s, annealing at 60 °C for 30 s, and amplification at 72 °C for 45 s/1 kb. A final 5-10 min cycle at 72 °C was routinely performed to allow the complete extension phase to occur

### Imaging and embryo measurement

Zebrafish embryos were anesthetized using MS-222 (Sigma-Aldrich) and imaged on an M165FC fluorescent stereomicroscope (Leica) with a DFC310 FX camera (Leica). For measuring the length of embryos, images were taken at 2.5x magnification and embryo length quantified in the CorelDRAW graphic suite. For confocal microscopy, images were taken from anesthetized embryos using a Leica TCS SP8 AOBS upright confocal using a 20x 0.50 Plan Flurotar objective and processed using LAS X (Leica version 3.5.2.18963).

### Quantitation of 28S:18S ratios

Total RNA was run on a TapeStation 4200 (Agilent) according to manufacturers instructions. Total RNA with integrity values in excess of 9 were selected for 28S:18S quantitation

### ASO-knockdown of U8 and rRNA processing assays in fibroblasts

ASO-mediated depletion of U8 in control cells (HCT116, colon carcinoma) was performed as described previously (16). RNA extraction and pre-rRNA processing analysis of U8 depleted cells and patient-derived fibroblasts (LCC) was performed as described previously (16, 43). The ATTCC fibroblast control PCS-201-012 (primary dermal fibroblasts normal human adult) was used. Sequence of anti-U8 ASO used in depletion: mGmGmAmUm*UATCCCACCTG*mAmCmGmAmU *N* and mN are deoxynucleotide and 2’-*O*-methoxyethylribonucleotide, respectively. Phosphodiester backbones are phosphorothioates.

### *In vitro* transcription of U8 RNA variants

U8 DNA templates containing a T7 consensus sequence were PCR amplified from human or zebrafish genomic DNA (see Supplementary Table 2 for primer sequences) and purified after agarose gel electrophoresis using a QIAEX II kit (Qiagen). Human and zebrafish U8 snoRNAs were generated using 400-1000ng of template DNA and a mMESSAGE mMACHINE T7 kit (Life Technologies) followed by Lithium Chloride precipitation and quantitation using a NanoDrop (Thermo Fisher Scientific)

### Microinjection and transgenesis

The yolk of fertilised 1-cell stage embryos were microinjected with 2nl of synthetic mRNA/snoRNA, or for transgenesis, with 40pg of both DNA and tol2 transposase mRNA (44), using a PLI-90 pico-injector (Harvard Apparatus) and Leica MZ6 stereomicroscope

### Electrophoretic mobility shift assays

For EMSAs, recombinant His-15.5K was incubated with 50,000 dpm ^32^P-end-labeled U8 snoRNA in EMSA buffer (20 mM HEPES-KOH, 150 mM KCl, 1.5 mM MgCl_2_, 0.2 mM EDTA and 0.1% Triton X-100) for 30–45 min on ice. Resulting RNA–protein complexes were resolved on a native 7% acrylamide gel for 8.5 h at 4 °C. Gels were dried and exposed to X-ray film for approximately 8 hr at −80 °C in the presence of an intensifying screen.

### 3′-processing assays

For 3′-processing assays, ^32^P-end-labeled U8 snoRNA was incubated with HeLa nuclear extract (CIL Biotech) at 30°C in buffer containing 0.25 mM ATP, 10 mM phosphocreatine, 3.2 mM MgCl_2_, 20 mM HEPES KOH, pH 7.9, 2.6% polyvinyl alcohol and 240 U RNasin (Promega). At 0, 30 and 60 min, 10 μl of the reaction was removed and added to a tube containing 4 μl of stop solution (1 mg/ml proteinase K, 50 mM EDTA and 1% SDS). Reactions were then incubated at 37 °C for 15 min, phenol extracted, precipitated and resolved on a 5% acrylamide/7 M urea gel. Gels were dried and exposed to X-ray film at −80 °C in the presence of an intensifying screen.

### Statistics and reproducibility

All statistical analyses were performed using GraphPad Prism 7.03 or Microsoft Excel software. Results are presented as mean ± s.d. For all analyses *P* < 0.05 was considered statistically significant. Statistical methods were not used to predetermine sample size, which varies between experiments. Experiments were not randomized. The investigators were not blinded to allocation during experiments and outcome assessment. For Fig. 1B significance was determined by one-way ANOVA and post-hoc Tukey’s multiple comparisons test. For fig. S3 significance was determined by a Mantel-Cox test. For all other statistical analyses’ significance was determined by an unpaired t-test. The number of biological replicates upon which significance was determined are specified in the figure legend

### Oligonucleotides used in this study

**Refer to** Table S2

## Acknowledgments

We thank the following people for reagents: pDest-γcrystallin:mCherry, pcs2^+^-mKate2 and pcs2^+^-H2B-mCerulean3 plasmids were gifts from Dr. Emily Don, pcs2^+^-nCas9n (Addgene plasmid #47929) was a gift from Dr. Adam Hurlstone. We thank Dr. Martin Reijns (Edinburgh) for helpful discussions. This study was supported by a grant to Y.J.C and R.T.O. from the Great Ormond Street Hospital Charity (V4017). P.R.K was supported by the Stroke Association (TSA LECT 2017/02). Research in the Lab of D.L.J.L. is supported by the Belgian Fonds de la Recherche Scientifique (F.R.S./FNRS), the Université Libre de Bruxelles (ULB), the Région Wallonne (DGO6) [grant RIBO*cancer* n°1810070], the Fonds Jean Brachet, and the International Brachet Stiftung.

## Author contributions

Conceptualization and design: A.P.B., Y.J.C., R.T.O., C.U., E.M.J., D.L.J.L.

Investigation: A.P.B performed the majority of the experiments and analysis of data with the exception of 2E, S5 (L.W, C.U.), 5B, 5C, S7, S9 (R.T.O.), 6C (S.C.) and Table S1 (G.I.R.)

Writing original draft: A.P.B., Y.J.C., R.T.O.

Writing – review and editing: all authors

---

**Supplementary Table S1.**
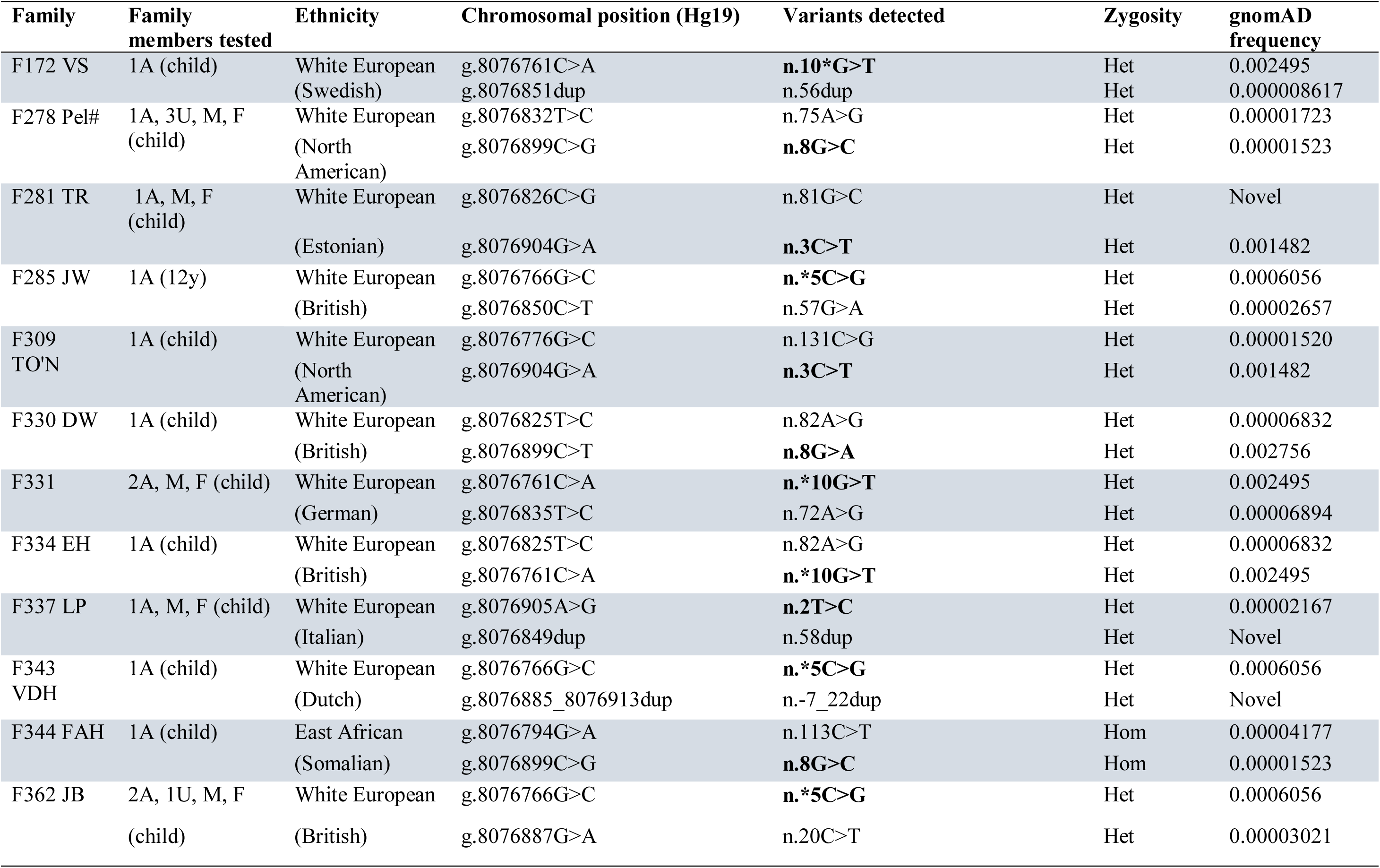
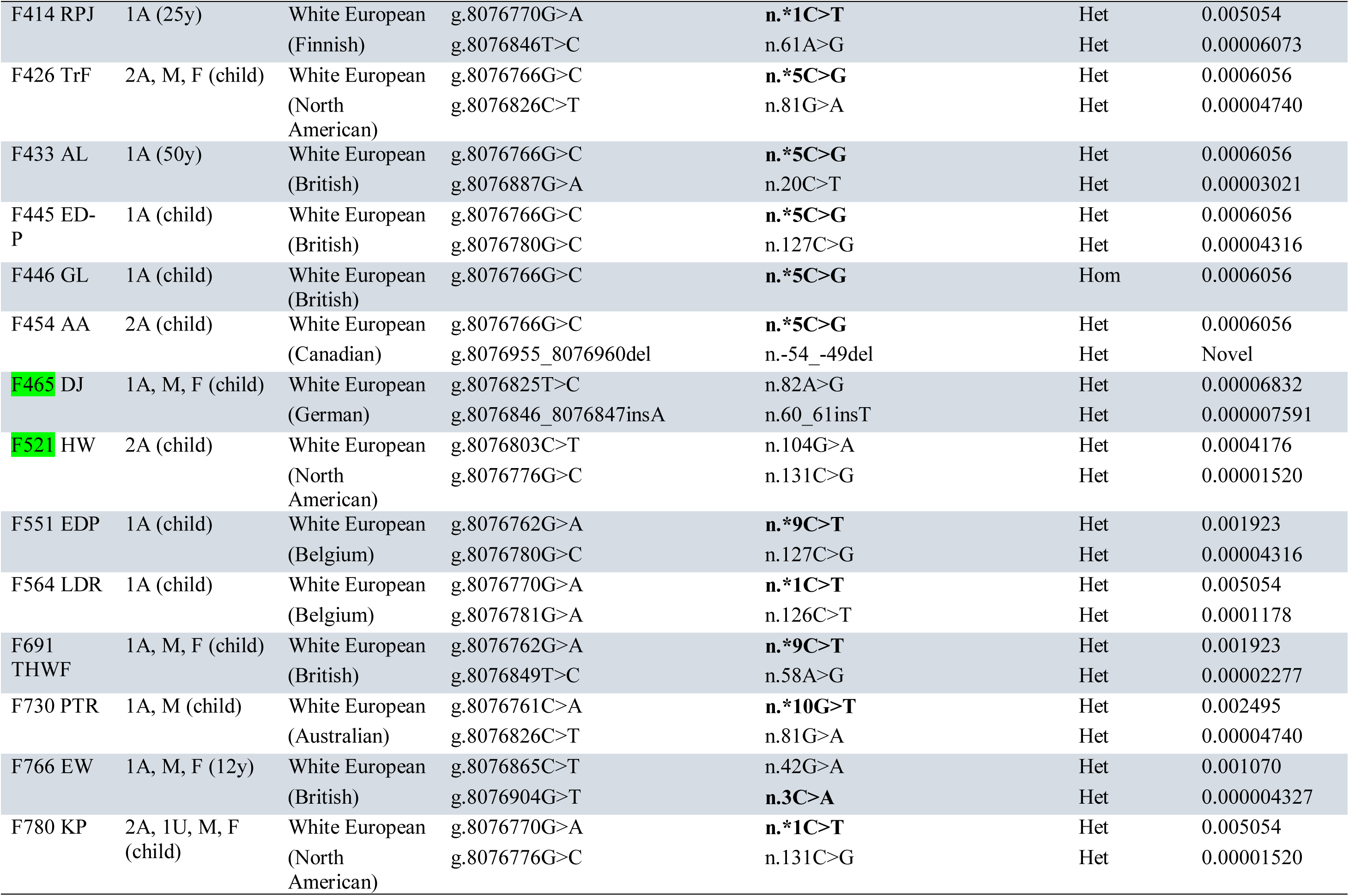
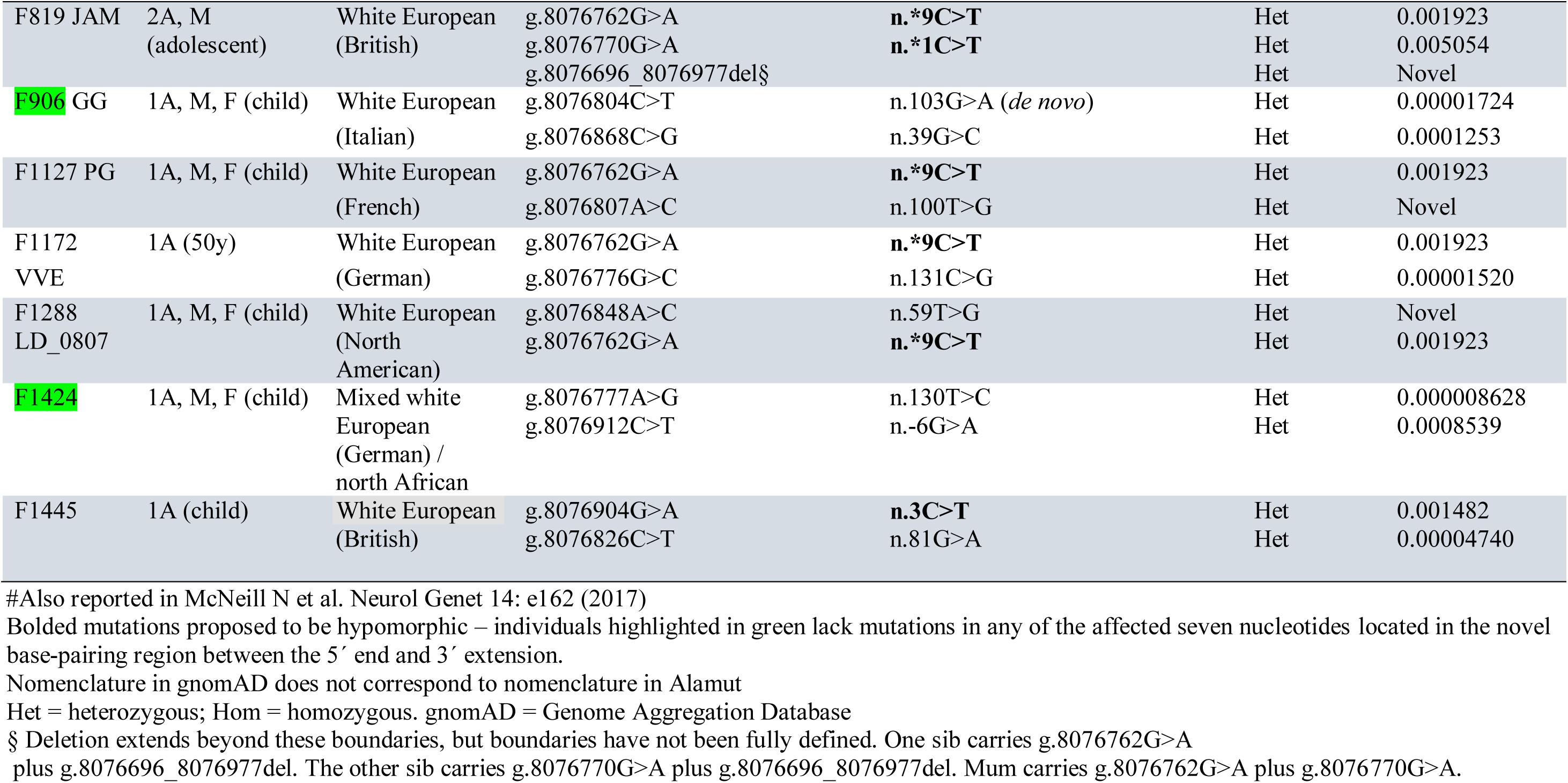
LCC cohort published in Jenkinson 2016.

**Supplementary Table S2.**
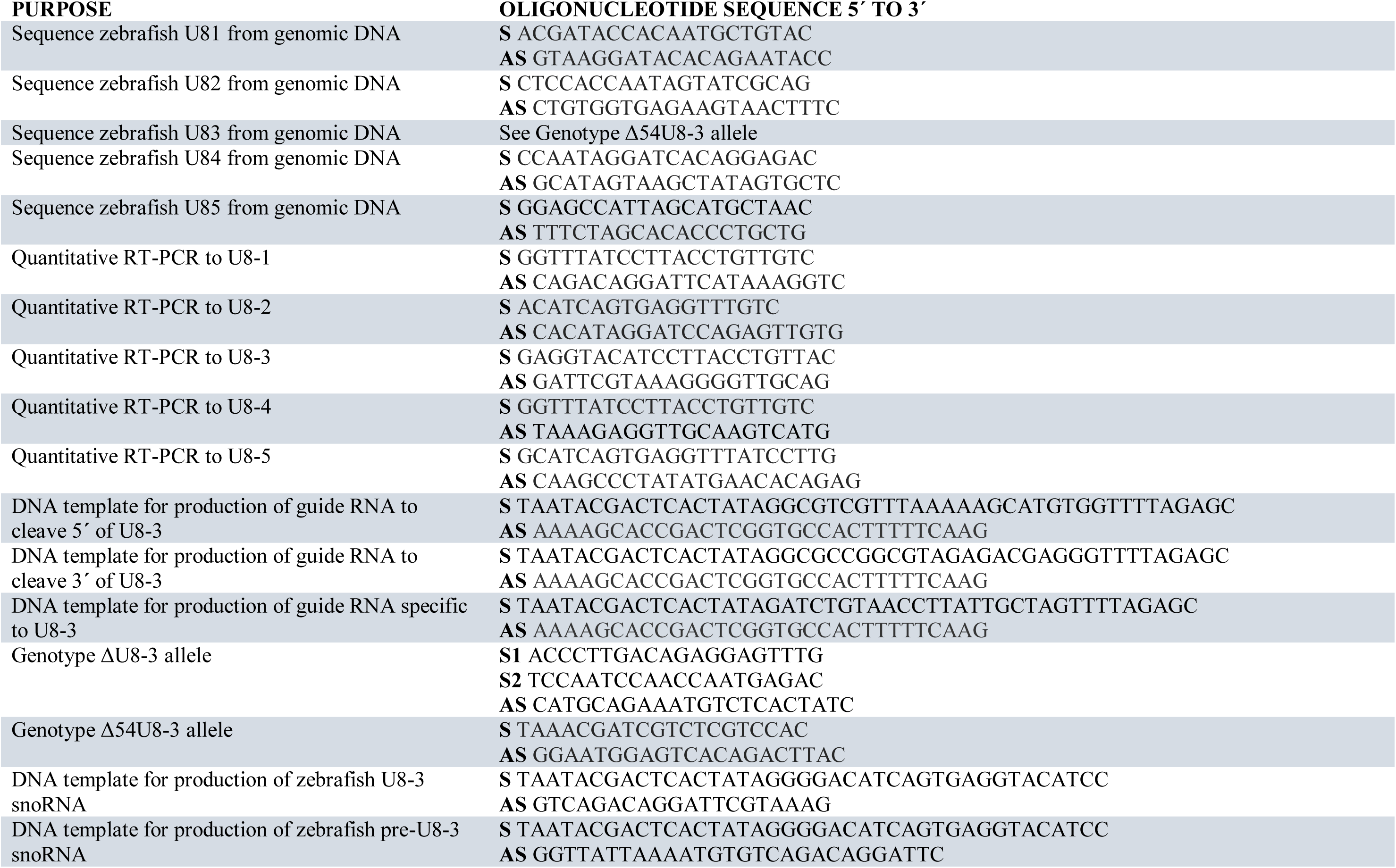
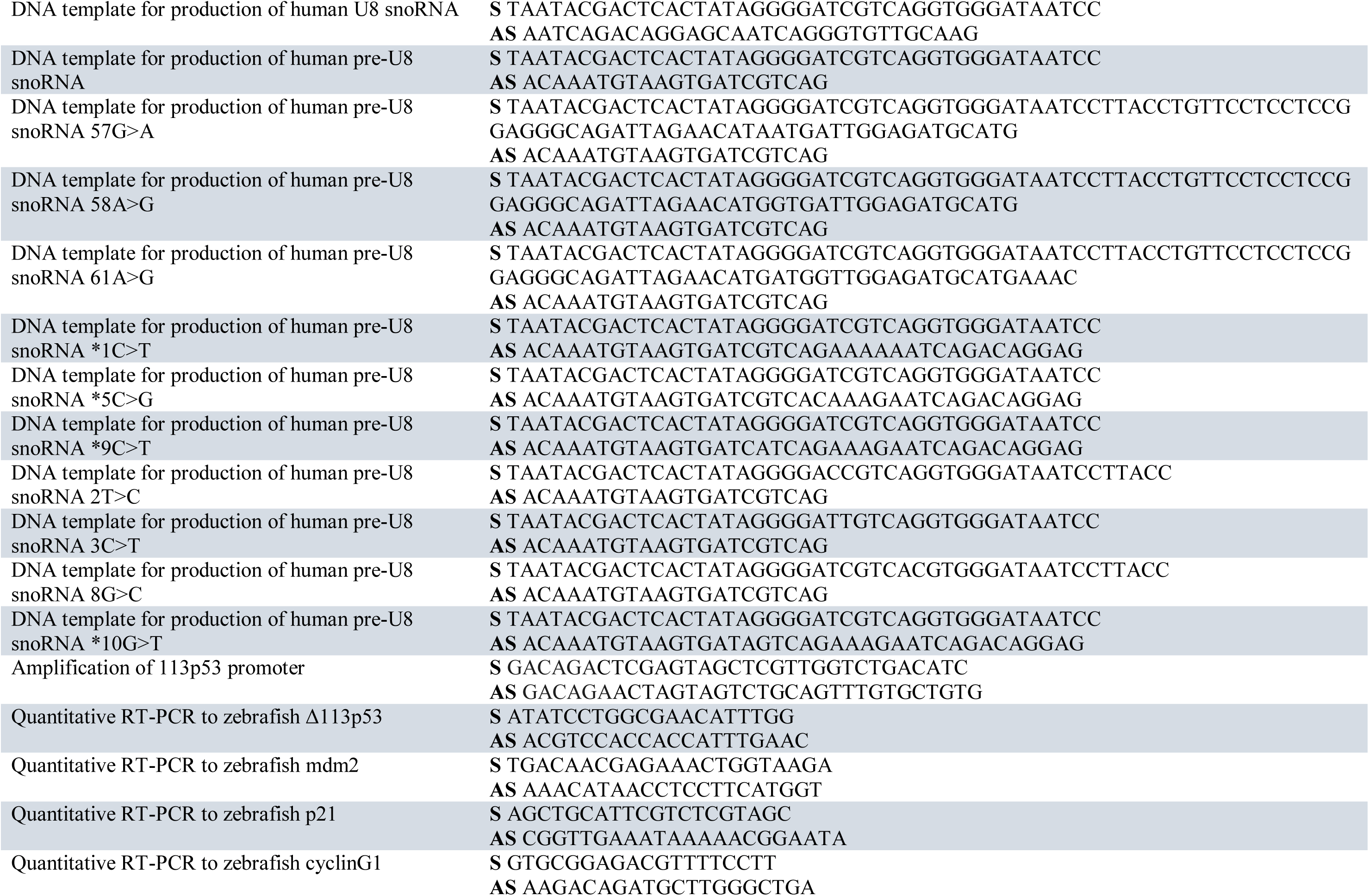
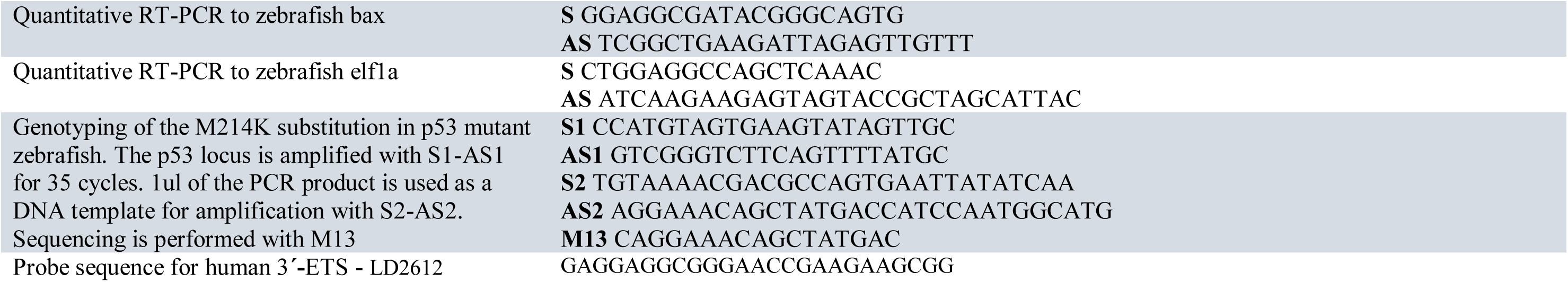
Oligonucleotides used in this study.

## Supplementary Figures

**Fig. S1.**
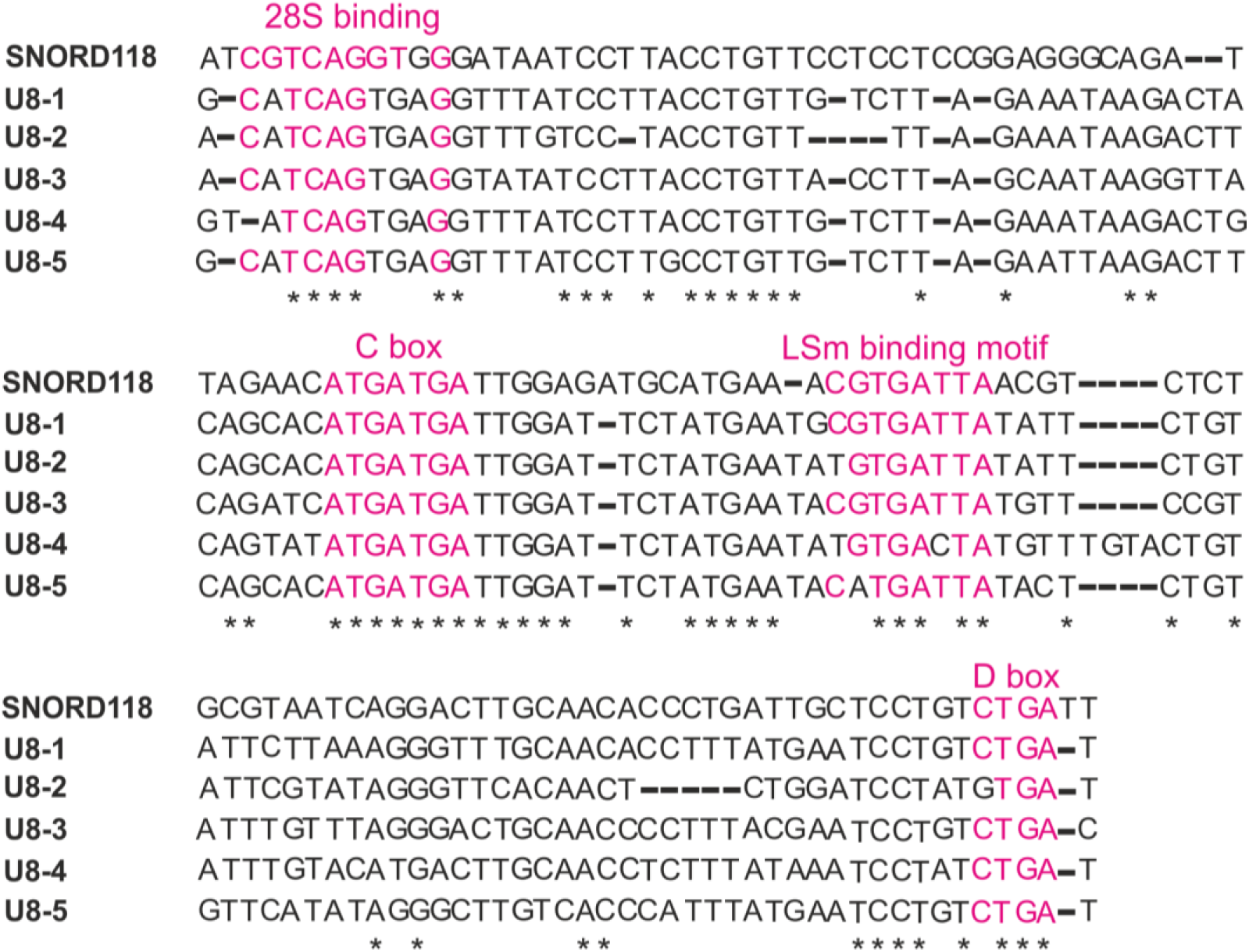
Human U8 and zebrafish U8 1-5 mature sequences exhibit extensive conservation. Mature human U8 and zebrafish U8 1-5 sequences were aligned using ClustalW (https://www.ebi.ac.uk/Tools/msa/tcoffee/). * denotes conserved nucleotides.

**Fig. S2.**
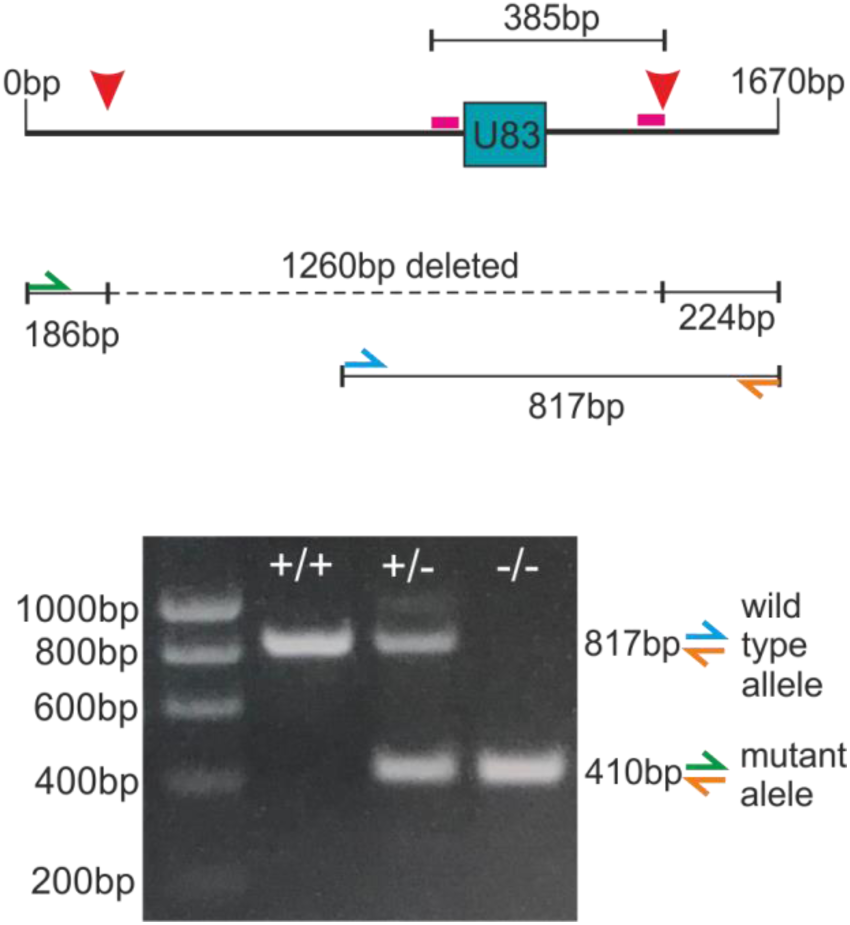
The ΔU8-3 mutant is homozygous for a 1.26kb deletion encompassing the *U8-3* gene locus. Schematic depicting the genotyping strategy for the zebrafish U8-3 mutant. Two sgRNAs predicted to excise 385bp encompassing the *U8-3* gene are represented by pink bars. Red arrowheads depict actual excision sites that delete 1260bp (dashed line) encompassing the *U8-3* gene locus. A three primer PCR was performed to genotype whereby 817bp (blue, orange primer) is amplified in the presence of a wildtype copy of *U8-3* and 410bp (green, orange primer) amplified in the presence of a deleted *ΔU8-3* allele. bp, base pairs.

**Fig. S3.**
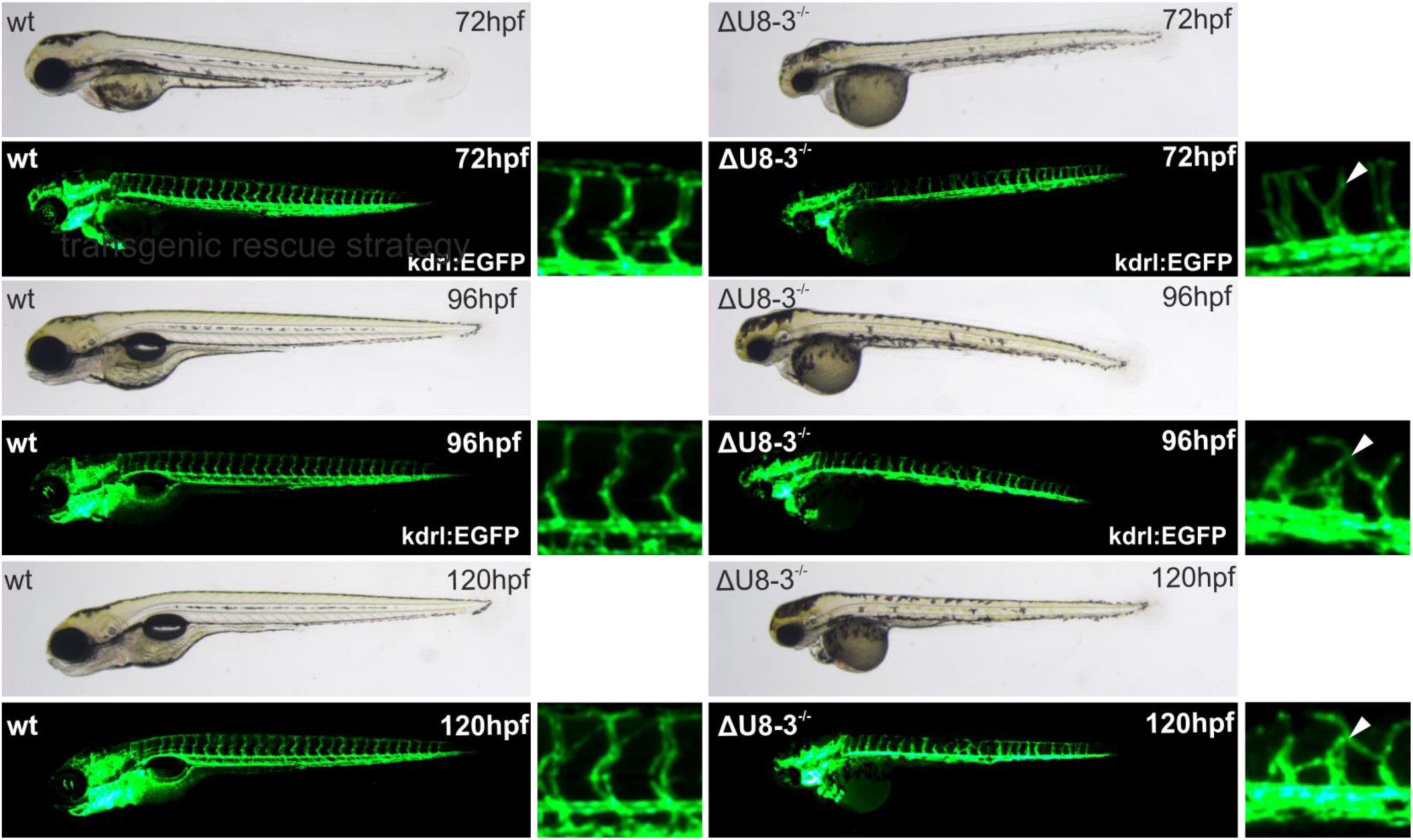
The ΔU8-3 mutant exhibits a range of gross morphological abnormalities and comes to a developmental standstill. From 72hpf ΔU8-3 mutants display impaired yolk resorption, reduced craniofacial structures and hindbrain swelling. By 96hpf ΔU8-3 mutants fail to inflate their swim bladders, display an underdeveloped intestine, and by 120hpf cardiac oedema. At all stages disorganized trunk vasculature is observed (white arrowheads) when compared to wildtype.

**Fig. S4.**
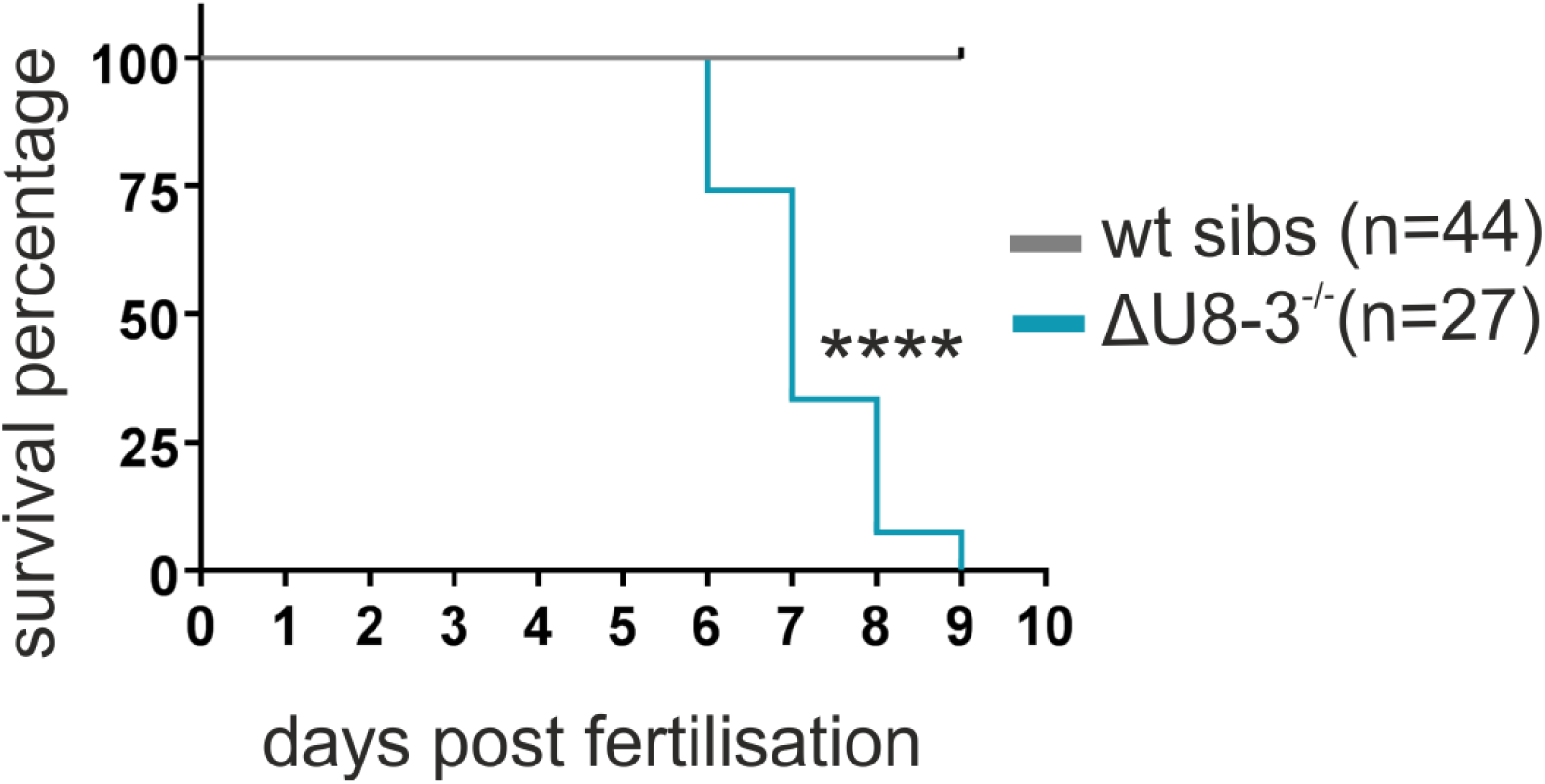
The ΔU8-3 mutant phenotype is lethal. Kaplain-Meier plot demonstrating that ΔU8-3 mutants begin to die from 6dpf and exhibit 100% mortality by 9dpf. dpf-days post fertilization.

**Fig. S5.**
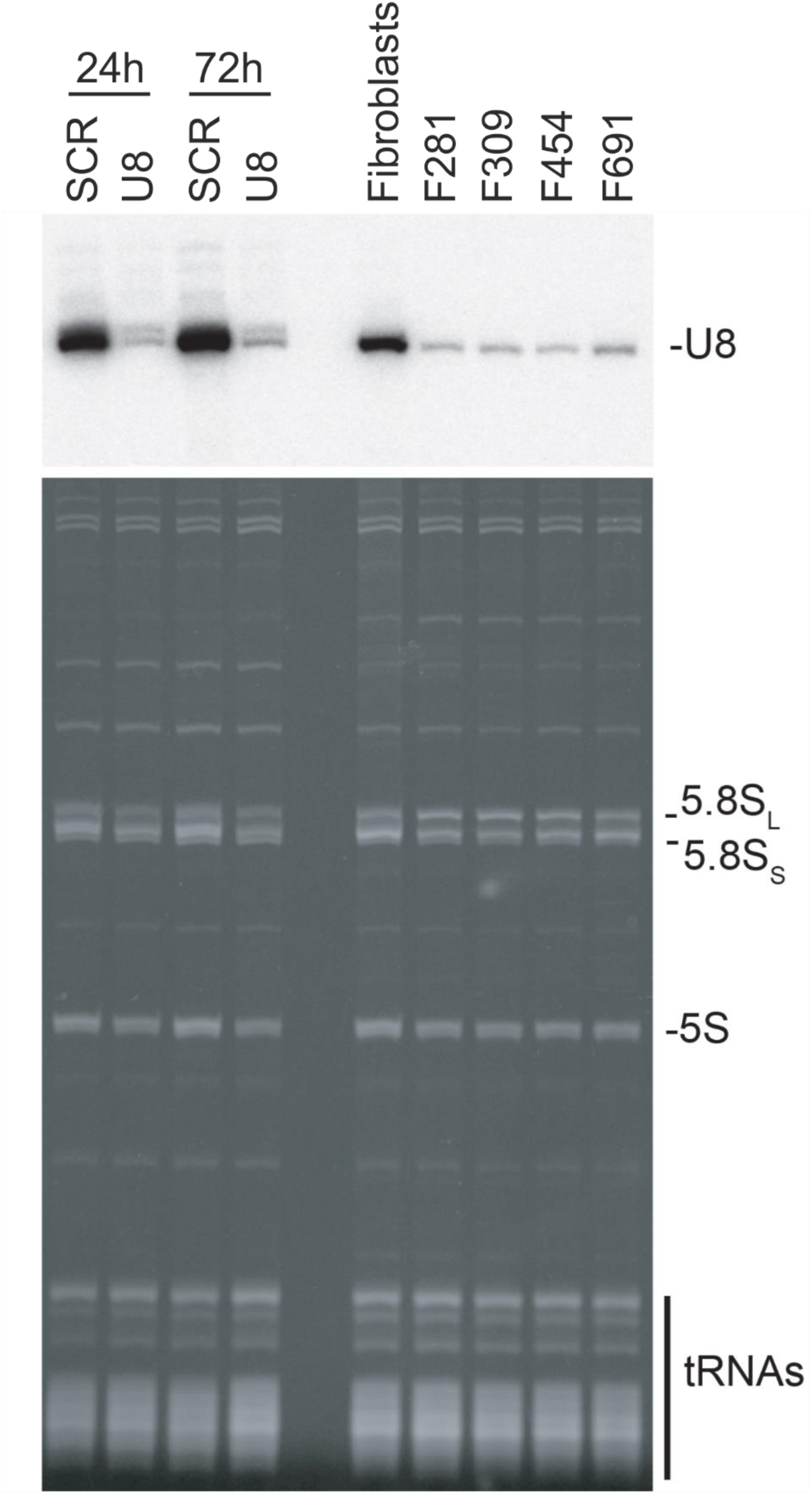
LCC patient fibroblasts express less total U8 compared to wildtype fibroblasts. Northern blot analysis with a probe specific to U8 snoRNA. U8 expression is shown after ASO-mediated depletion of human U8 snoRNA from HCT116 cells in which rRNA processing was analyzed (see Figure 2E), or in LCC patient fibroblast cell lines F281, F306, F454 and F691. ASO – antisense oligonucleotide. The same amount of total RNA (OD260) was loaded in each lane of the gel. Ethidium bromide staining of the acrylamide gel demonstrates equal loading, as determined by tRNAs which are produced independently.

**Fig. S6.**
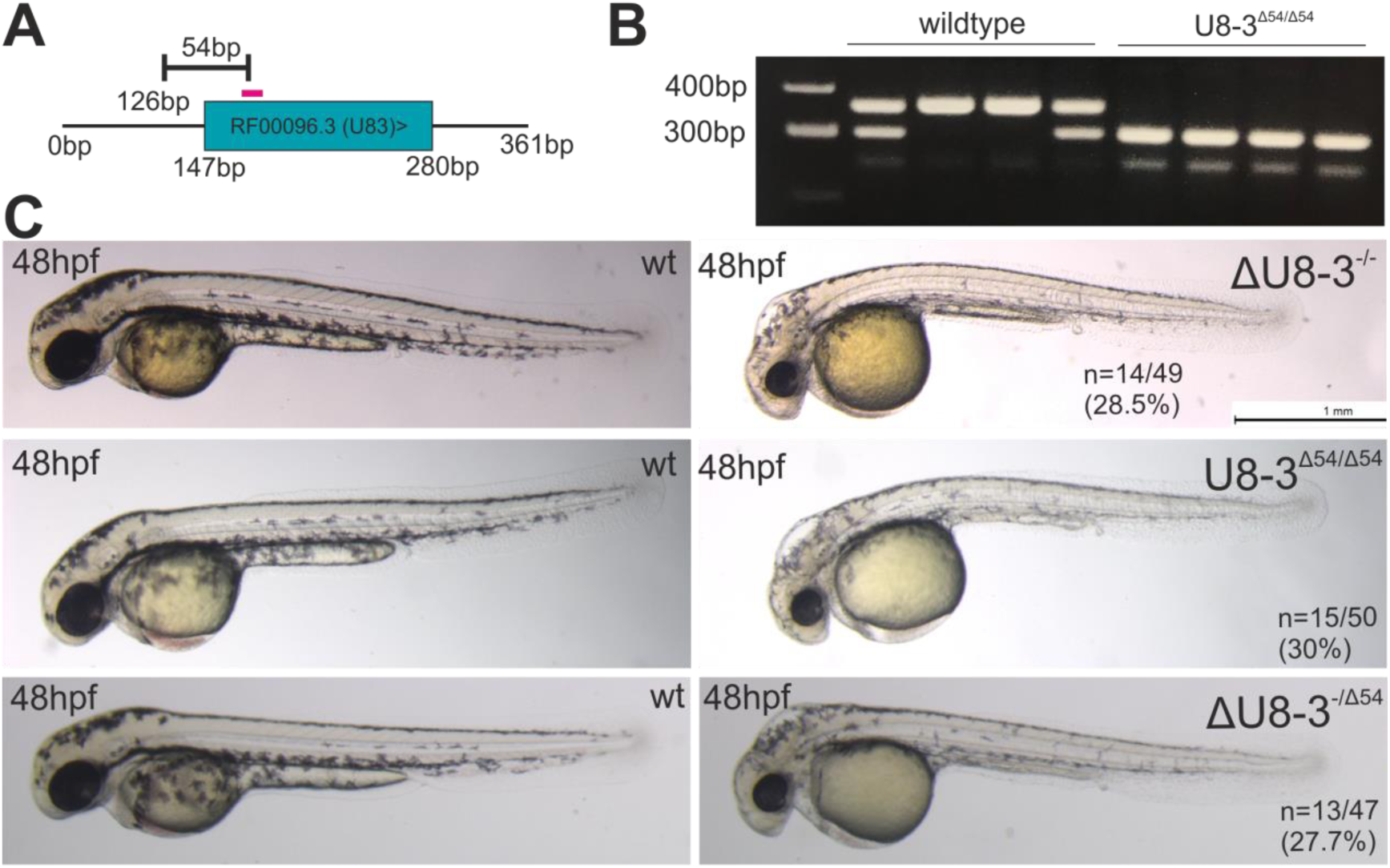
Non-complementation with independent mutant alleles of *U8-3* demonstrates that the ΔU8-3 mutant is specific. **A)** Schematic depicting the 54bp deletion allele (Δ54U8-3) generated with a sgRNA (pink bar) from the sgRNAs used to excise *U8-3* shown in Figure 2A. **B)** Genotyping demonstrates the 54bp deletion (307bp band) segregates with the U8-3 mutant phenotype. **C)** Both ΔU8-3 and Δ54U8-3 alleles display mendelian recessive inheritance i.e. approximately 25% of mutant progeny are produced from heterozygous carriers. When heterozygous carriers of each allele were crossed to each other non-complementation (failure to produce 100% wildtype progeny) was observed, demonstrating the alleles result from loss-of-function in the same gene.

**Fig. S7.**
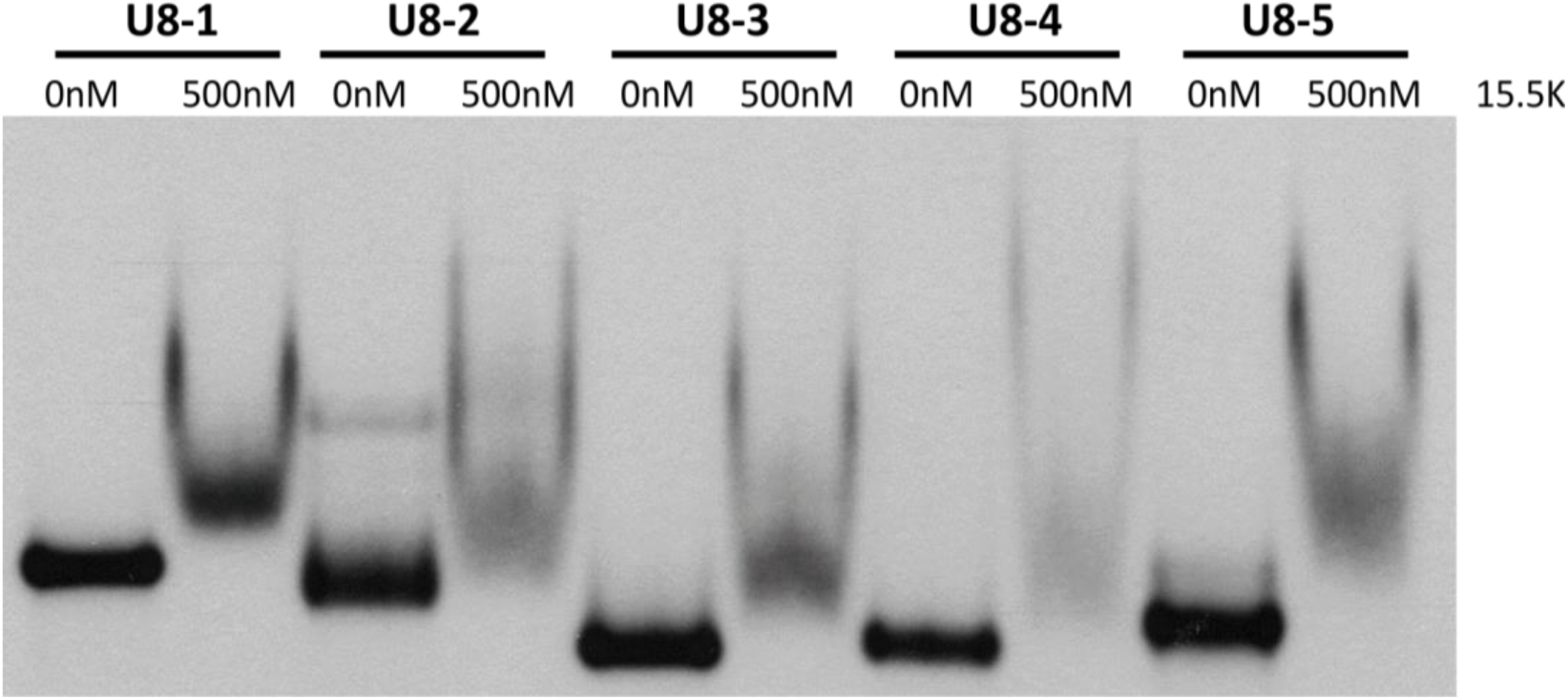
EMSAs demonstrates all five zebrafish U8s bind 15.5K. Protein binding of zebrafish U8 variants. Electrophoretic mobility shift assays (EMSAs) using 5’-end-radiolabeled *in vitro* transcribed mature zebrafish *U8-1*, *U8-2*, *U8-3*, *U8-4* and *U8-5* with 500nM of His6-tagged 15.5K human protein (15.5K). Binding of 15.5K to zebrafish U8 is indicated by a mobility shift.

**Fig. S8.**
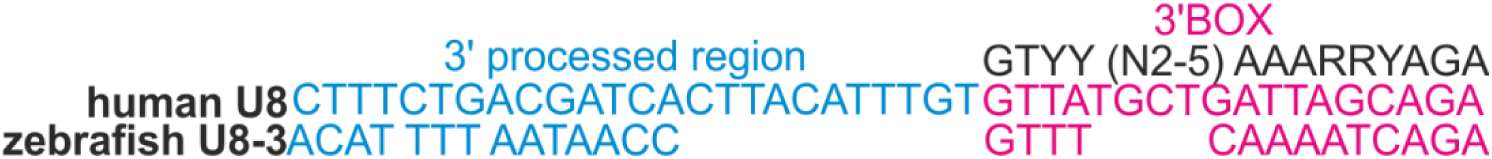
Alignment of human U8 and zebrafish 3’ extension sequences. Human and putative zebrafish 3’ extension sequences (blue) are divergent. A 3’ BOX is predicted for zebrafish U8-3 (pink) according to the 3’ BOX consensus sequence (black).

**Fig. S9.**
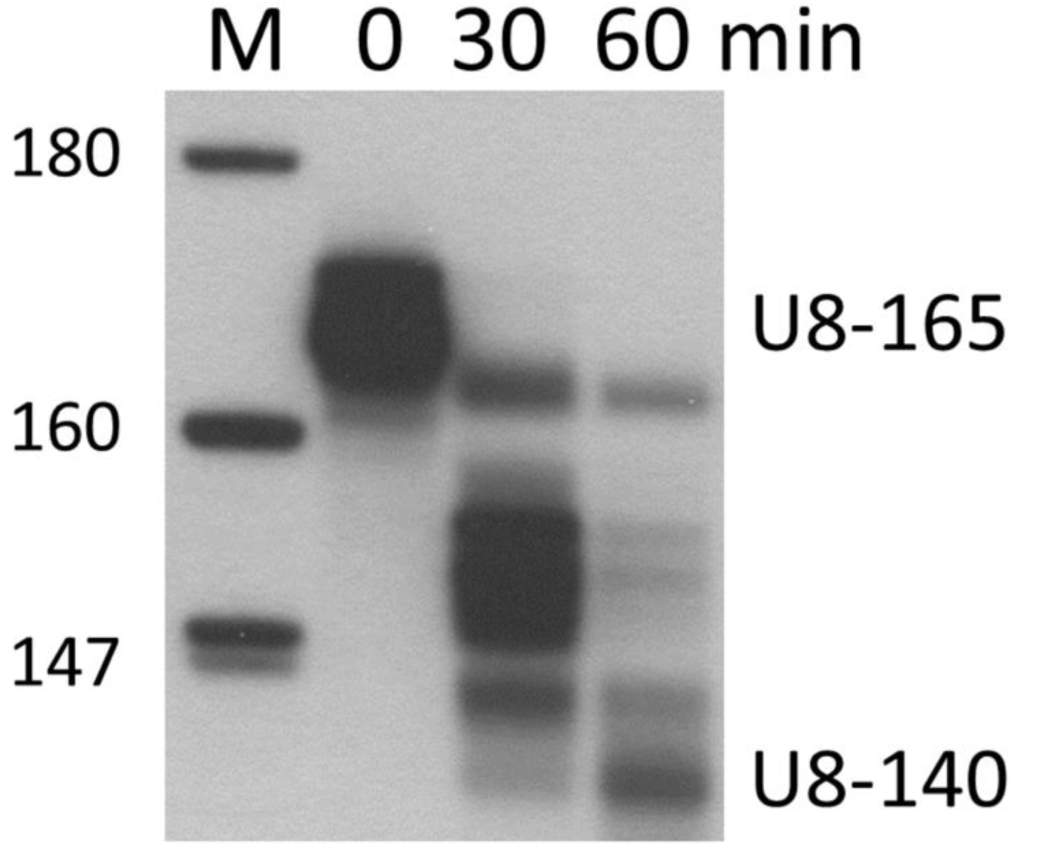
Time course of wild type human pre-U8 snoRNA processing. Processing of 5’ end-radiolabelled *in vitro* transcribed precursor U8 wildtype snoRNA (U8-165) was assessed in HeLa nuclear extracts at 0, 30 and 60 minutes (min) to monitor production of mature U8 (U8-140). Mature U8 is clearly present after 60 min of processing.

**Fig. S10.**
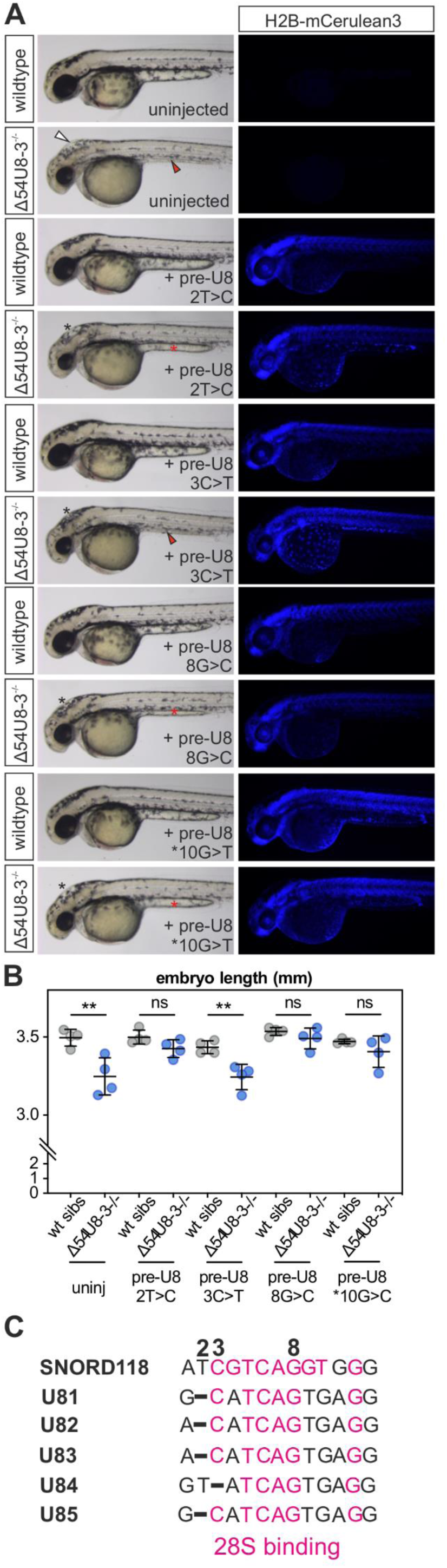
Functional testing of four LCC patient mutant U8 snoRNAs in zebrafish. **A)** Representative images of rescue experiments conducted with four putatively hypomorphic U8 mutations detected in LCC patients. White arrowhead indicates hindbrain swelling and red arrowheads indicate abnormal yolk extension. Black and red asterisks indicate rescued hindbrain swelling and yolk extension respectively **B)** pre-U8^3C>T^ does not rescue the reduced embryo length of ΔU8-3 mutants, whereas pre-U8^2T>C^, pre-U8^8G>C^ and pre-U8^*10G>T^ restores ΔU8-3 mutant embryo length to that of wildtype siblings. n=4 embryos per genotype **C)** alignment of human U8 and zebrafish U8 28S interaction region. Shared identity between zebrafish and human 28S-binding nucleotides is shown in pink

**Fig. S11.**
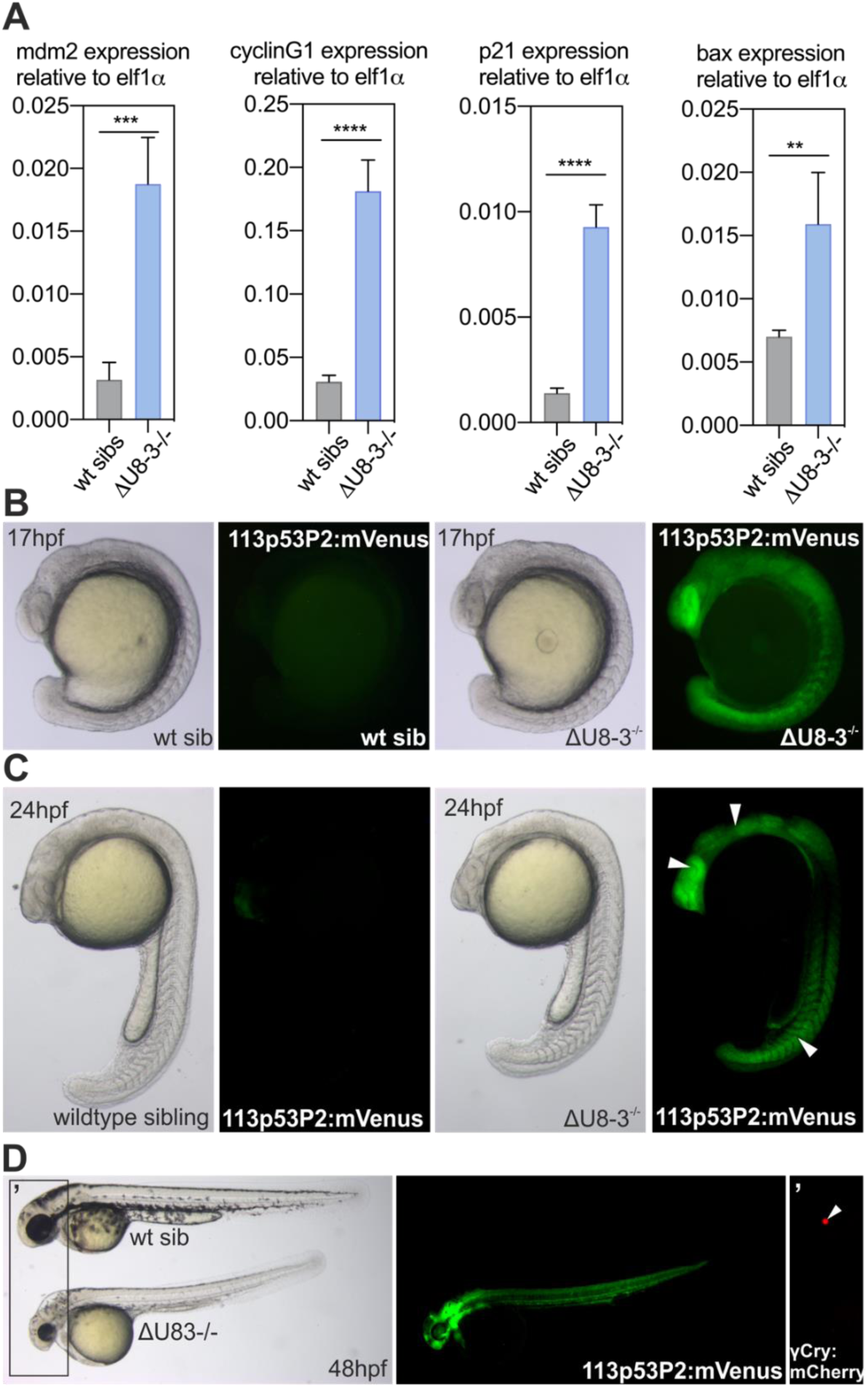
tp53 is trans-activated in response to loss of *U8-3* in zebrafish. **A)** quantitative RT-PCR demonstrates significant upregulation of the tp53 target genes *mdm2*, *cyclinG1*, *p21* and *bax* in ΔU8-3 mutants when compared to wildtype siblings at 24hpf. **B)** tp53 activity is observed throughout the U8-3 mutant at 17hpf, prior to the onset of observable gross morphological abnormalities, particularly in the eye. **C)** tp53 activity is observed in numerous tissues that develop abnormally in ΔU8-3 mutants, including the eye, hind-brain and somites (arrowheads) at 24hpf. **D)** tp53 activity is observed throughout the ΔU8-3 mutant at 48hpf. Integrated 113p53P2:mVenus transgene is reported by γ-crystallin:mCherry in the backbone of the plasmid which drives lens specific mCherry expression from approximately 36hpf. The developmental delay in ΔU8-3 mutants delays expression of this reporter.

**Fig. S12.**
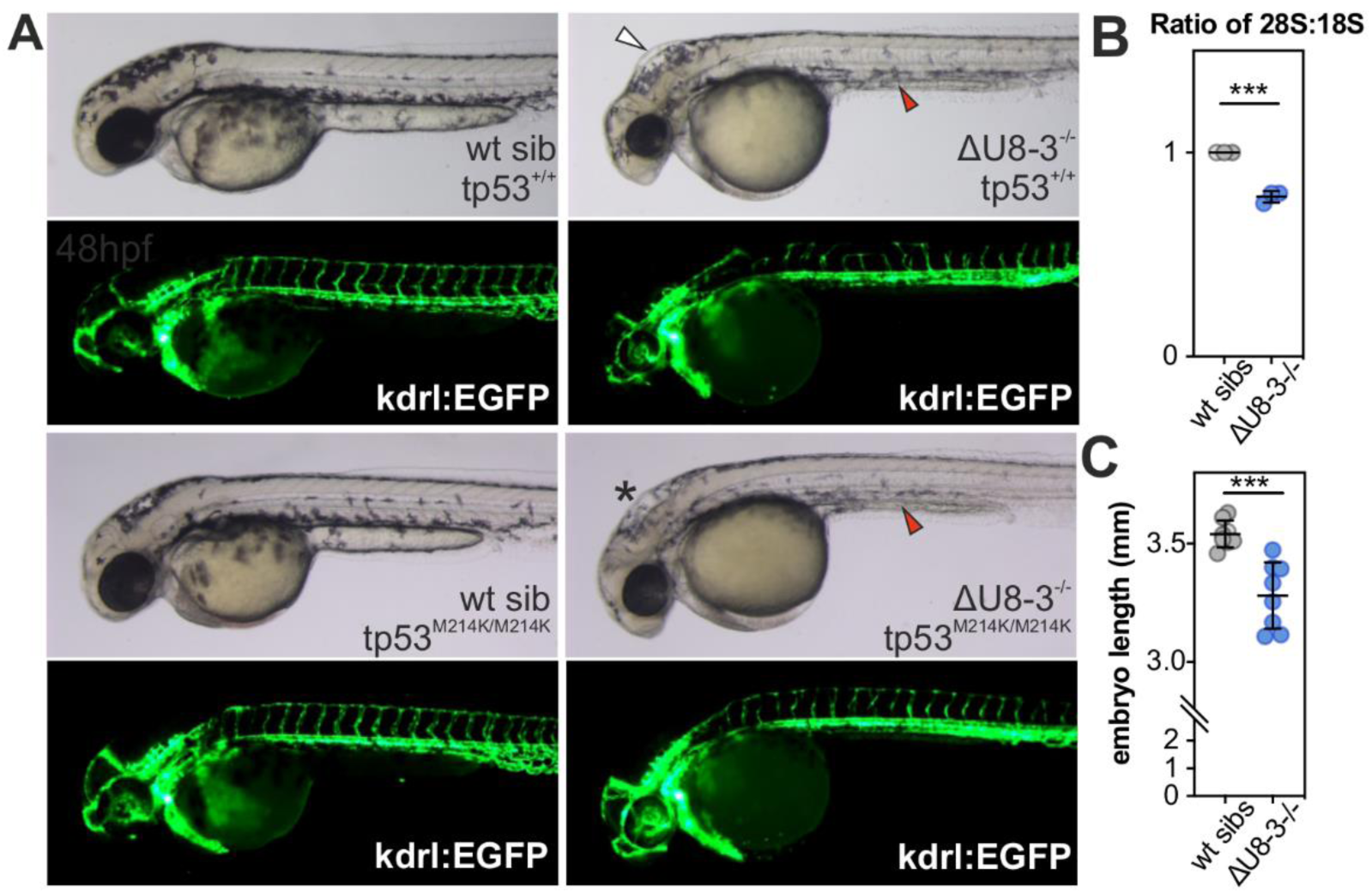
Inactivation of tp53 partially rescues the ΔU8-3 mutant phenotype though fails to rescue the embryo length or rRNA biogenesis defect. **A)** representative images showing inactivation of tp53 signaling rescues the hindbrain swelling of ΔU8-3 mutants (white arrowhead compared to black asterisk) and the trunk vasculature, but not the yolk extension (red arrowhead) abnormality. **B)** Tapestation analysis demonstrates tp53 mutant ΔU8-3 mutant embryos display a preferential reduction in 28S biogenesis at 48hpf. n=3 biological replicates per genotype. **C)** tp53 mutant U83 mutant embryos are significantly shorter than tp53 mutant U8-3 wildtype sibling embryos at 48hpf. n=8 biological replicates per genotype.

